# RNA elements required for the high efficiency of West Nile Virus-induced ribosomal frameshifting

**DOI:** 10.1101/2024.10.16.618579

**Authors:** Nikolay A. Aleksashin, Conner J. Langeberg, Rohan R. Shelke, Tianhao Yin, Jamie H. D. Cate

**Author notes:** To whom correspondence should be addressed: Jamie H. D. Cate, Innovative Genomics Institute, 2151 Berkeley Way, Rm. 212C, Berkeley, CA, USA, 94720.

## Abstract

West Nile Virus (WNV), a member of the *Flaviviridae* family, requires programmed -1 ribosomal frameshifting (PRF) for translation of the viral genome. The efficiency of WNV frameshifting is among the highest observed to date. Despite structural similarities to frameshifting sites in other viruses, it remains unclear why WNV exhibits such a high frameshifting efficiency. Here we employed dual-luciferase reporter assays in multiple human cell lines to probe the RNA requirements for highly efficient frameshifting by the WNV genome. We find that both the sequence and structure of a predicted RNA pseudoknot downstream of the slippery sequence–the codons in the genome on which frameshifting occurs–are required for efficient frameshifting. We also show that multiple proposed RNA secondary structures downstream of the slippery sequence are inconsistent with efficient frameshifting. We mapped the most favorable distance between the slippery site and the pseudoknot essential for optimal frameshifting, and found the base of the pseudoknot structure likely is unfolded prior to frameshifting. Finally, we find that many mutations in the WNV slippery sequence allow efficient frameshifting, but often result in aberrant shifting into other reading frames. Mutations in the slippery sequence also support a model in which frameshifting occurs concurrent with or after translocation of the mRNA and tRNA on the ribosome. These results provide a comprehensive analysis of the molecular determinants of WNV-programmed ribosomal frameshifting and provide a foundation for the development of new antiviral strategies targeting viral gene expression.

## Introduction

Programmed ribosomal frameshifting (PRF) is a translational recoding mechanism that allows viruses and other organisms to regulate gene expression by altering the mRNA reading frame of ribosomes during translation. This process is especially prevalent in RNA viruses, where frameshifting enables the production of multiple proteins from a single mRNA, thereby increasing the coding capacity of the viral genome (1, 2). Among the many viruses that employ PRF, West Nile Virus (WNV) is a member of the *Flaviviridae* family, an arthropod-borne virus primarily transmitted through the bite of infected mosquitoes, particularly those of the *Culex* species (3, 4). First identified in the West Nile district of Uganda in 1937 (5), WNV has since spread globally, becoming a significant public health concern (6). The virus is known to infect a wide range of hosts, including birds, which are its primary reservoir, and mammals, including humans and horses (3, 7). In humans, WNV infections are often asymptomatic, but in some cases, they can lead to severe neurological diseases such as encephalitis, meningitis, and hemorrhagic fever (8–10). To date, no preventive vaccines or targeted treatments exist for WNV infection (11).

WNV is a positive-sense single-stranded RNA virus with a genome approximately 11,000 nucleotides (nts) in length (12, 13) (**Figure 1A**). It encodes a single polyprotein that is co- and post-translationally processed by host and viral serine proteases (14) into three structural proteins (C, prM, and E) and seven nonstructural proteins (NS1, NS2A, NS2B, NS3, NS4A, NS4B, and NS5) (15, 16). The NS proteins are primarily involved in viral replication and assembly, with NS5 serving as the RNA-dependent RNA polymerase and NS3 as a protease and helicase (13, 16, 17). While the molecular function of NS1 is not fully understood, its expression is unique in WNV and some other flaviviruses because it exists in two forms: full-length NS1 and an extended NS1′ (18–20). The production of NS1′ is dependent on a -1 ribosomal frameshift that occurs within the NS2A gene, leading to the continued translation in a -1 reading frame that harbors a premature stop codon (15, 21–23). While NS1 plays a critical role in viral replication and assembly, NS1′ is not essential. However, NS1′ contributes to WNV virulence. Studies have shown that the absence of NS1′ reduces neuroinvasiveness (22), viral replication (23), and viral RNA levels (24). Inhibition of frameshifting decreases the ratio of structural to nonstructural proteins, such as E/NS5, and results in reduced virus production (23). Furthermore, mutations that attenuate -1 PRF have been associated with lower virulence in mouse models of encephalitis (22). A WNV clone deficient in -1 PRF exhibited reduced infectivity in birds, while a WNV with impaired frameshifting showed decreased replication and spread in *Culex* mosquitoes (23). Thus, targeting the WNV -1 PRF could present a potential therapeutic approach for combating WNV infections (25).

**Figure 1.**
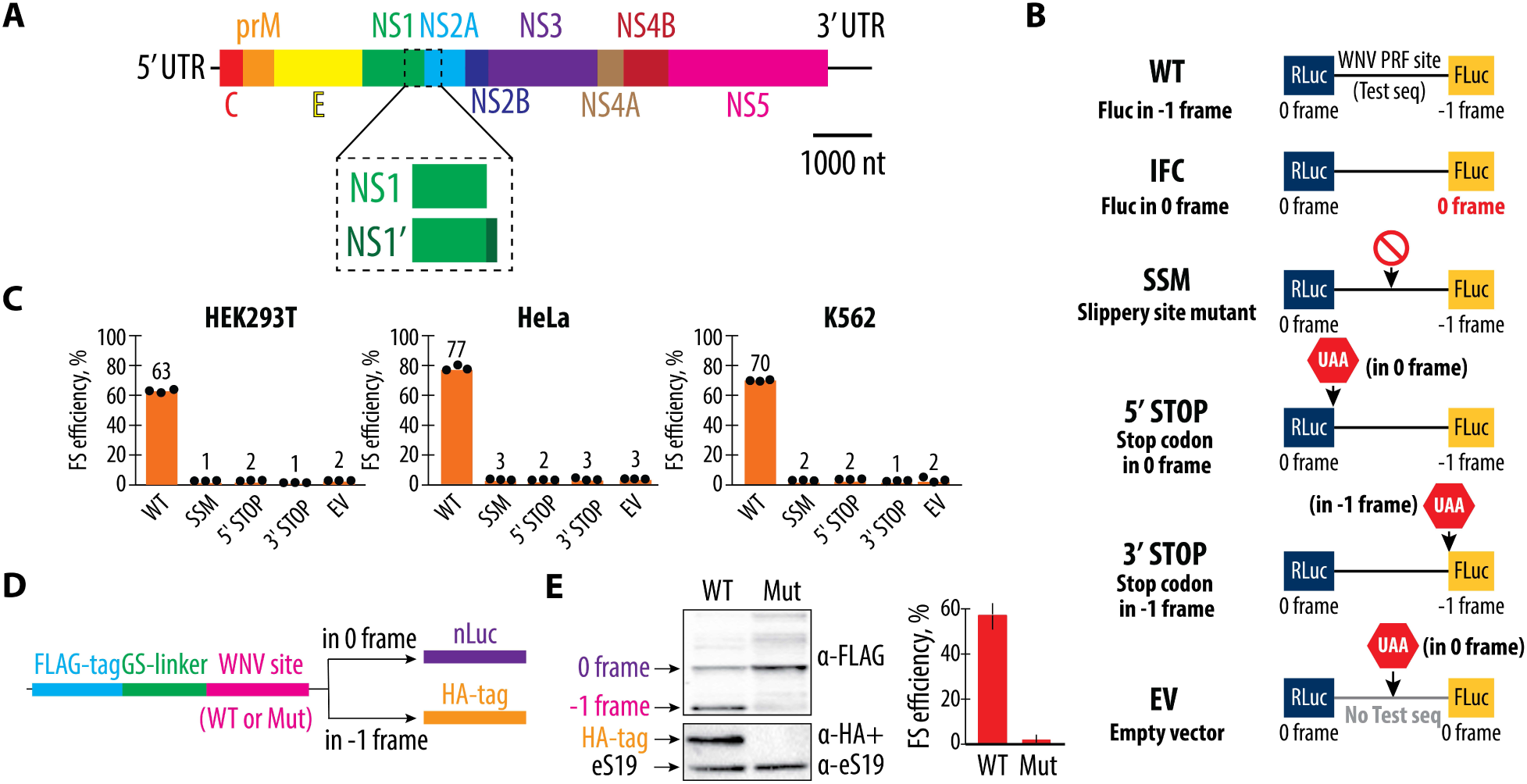
Programmed -1 ribosomal frameshifting (PRF) at the NS1 and NS2A region of the West Nile Virus (WNV) genome. (**A**) Schematic representation of the WNV genome. The genome consists of a single positive-sense RNA that encodes a polyprotein, which is co- and post-translationally processed into three structural proteins (C, prM, and E) and seven nonstructural proteins (NS1, NS2A, NS2B, NS3, NS4A, NS4B, and NS5). The NS1 and NS2A junction includes the programmed -1 ribosomal frameshifting site, which leads to the production of both the full-length NS1 and the extended NS1′ protein. (**B**) Diagram of the dual-luciferase reporter constructs used to measure frameshifting efficiency. The wild-type (WT) reporter contains Renilla luciferase (RLuc) in the reference reading frame (0 frame) and firefly luciferase (FLuc) in the -1 reading frame, separated by the WNV PRF site. The IFC construct controls for no frameshifting by placing FLuc in the 0 frame. The mutations in the slippery site (SSM) are designed to abolish frameshifting. The 5′ STOP and 3′ STOP constructs introduce stop codons in either the 0 or -1 frames before or after the testing sequence, respectively, while the empty vector (EV) contains no test sequence. See the Materials and Methods for more on the use of the IFC construct for normalizing experiments. (**C**) The -1 frameshifting efficiencies of the wild-type (WT) and mutant constructs in HEK293T, HeLa, and K562 cell lines. Frameshifting efficiency is shown as a percentage of the ratio between normalized FLuc/RLuc luminescence in the -1 frame to the sum of normalized Fluc/Rluc luminescence in -1 and 0 frames. Bars represent the means of three independent experiments, with each dot corresponding to the individual experimental data points. (**D**) Schematic of the reporter construct used in *in vitro* translation system based on human extracts (44). The reporters include either wild-type (WT) or mutant (Mut) WNV frameshifting sites. While the translation products always carry a 3x FLAG tag followed by GS-linker on their N-terminus, only -1 frameshifting product polypeptides have an HA tag on their C-terminus. Ribosomes that continue translation in the 0 frame synthesize nanoluciferase (nLuc). (**E**) Western blot analysis of FLAG and HA tags in WT and mutant constructs. The WT construct shows both FLAG- and HA-tagged proteins, indicating efficient frameshifting. The mutant (Mut) construct abolishes frameshifting, as indicated by the absence of the HA-tagged product, while increasing nanoluciferase protein levels as detected by the α-FLAG band. Quantification of frameshifting (FS) efficiency is shown on the right.

Despite advances in understanding PRF in other viruses (25–27), the exact mechanism and molecular determinants of the high frameshifting efficiency in WNV remained to be uncovered. The molecular mechanism underlying -1 PRF in WNV involves a highly conserved slippery sequence, a heptanucleotide motif (5’-X_XXY_YYZ-3’) that facilitates tRNA slippage, and a downstream RNA secondary structure, predicted to be a pseudoknot that might induce ribosomal pausing (28–30). The slippery sequence is the site where the ribosome shifts from one reading frame to another, while the downstream RNA structure is thought to act as a mechanical barrier, slowing down the ribosome and promoting the frameshifting event. In other viral systems, such as HIV, the frameshifting-stimulating element may adopt alternative structures, either forming a stem-loop or a pseudoknot, depending on specific RNA conditions (31–35). Recent single-molecule tweezer experiments suggest that the frameshifting-stimulating element in WNV could also form alternative structures, which may influence the efficiency of the frameshifting event (27). In WNV, the slippery sequence is usually 5’-C_CCU_UUU-3’, which is well-conserved among different strains of the virus (36, 37). In WNV, the specific contributions of the nascent peptide sequence and the surrounding mRNA regions in modulating WNV frameshifting efficiency are not known. Additionally, the sequence conservation of the frameshifting sites across different WNV strains and its implications for frameshifting efficiency have yet to be fully delineated. Here, we conducted a comprehensive analysis of WNV frameshifting. We first examined the role of the sequence spanning the NS1 and NS2A genes in promoting ribosomal frameshifting and identified the minimal elements required for this process, focusing on both the nascent peptide and the mRNA sequence. We then explored the structural integrity and sequence conservation of the predicted pseudoknot downstream of the frameshifting site, as well as the spatial relationship between the slippery site and the pseudoknot. We also investigated how mutations to the slippery sequence influence -1 frameshifting outcomes, as well as the induction of alternative frameshifting events such as -2 frameshifting. Taken together, these studies provide a foundation for understanding how WNV uses multiple RNA features of the -1 PRF to induce highly efficient frameshifting.

## Results

### Minimal sequence requirements for efficient West Nile Virus -1 programmed ribosomal frameshifting

To identify the minimal sequence requirements for efficient WNV -1 PRF, we first explored the role of sequences spanning the junction of the NS1 and NS2A genes. We employed a dual-luciferase reporter (38, 39) in three different human cell lines: HEK293T, HeLa, and K562. These cell lines were chosen to represent a range of biological contexts, ensuring that our findings are not cell-type specific. The reporter constructs include a wild-type (WT) sequence containing 258 nucleotides from the WNV genome (strain NY99, RefSeq NC_009942, nucleotides 3420-3678) (40), as well as several constructs designed to dissect the contributions of different sequence elements to frameshifting efficiency, as well as controls to ensure -1 PRF is properly detected (**Figure 1B**) (39). These controls include a slippery sequence mutant (SSM) designed to abolish frameshifting by disrupting the critical sequence where ribosomal slippage occurs, as well as constructs containing premature stop codons inserted either in the 0-frame or -1 frame (38, 41, 42). With WT sequences, we observed high levels of frameshifting across all cell lines, with the most pronounced effect in HeLa cells, where frameshifting reached up to 77% efficiency, in agreement with previous publications (25, 43) (**Figure 1C** and **Figure S1**). This robust frameshifting suggests that the 258-nucleotide segment of the WNV NS1 and NS2A genes contains all the necessary elements to effectively induce a -1 frameshift in a variety of cellular environments (33, 36, 37). As expected, mutating the slippery site abolished nearly all frameshifting, underscoring the essential role of the slippery site in this process (**Figure 1C**). Introduction of the stop codons upstream and downstream from the frameshifting site similarly nearly eliminated frameshifting, confirming that uninterrupted translation through both the 0-frame and the -1 frame is necessary for the successful production of the NS1′ protein.

The results from these dual-luciferase assays were corroborated by *in vitro* translation experiments using highly efficient translation extracts from HEK293T cells (44). These experiments allowed us to directly measure the production of FLAG- and HA-tagged proteins, which serve as markers for translation in the 0-frame and -1 frame, respectively (**Figure 1D**). The WT construct produced strong bands for both tags, indicating efficient frameshifting and successful translation in both reading frames. In contrast, the mutation that abolishes the slippery site sequence (SSM construct) resulted in a dramatic reduction in HA-tagged protein production, further validating the critical role of the NS2A gene’s specific sequences in promoting frameshifting (**Figure 1E**).

### Importance of the mRNA sequence but not nascent peptide sequence in WNV -1 frameshifting

Having identified the minimal mRNA requirements for WNV frameshifting, we next probed the molecular basis of the -1 PRF process. Frameshifting is a highly dynamic event that may depend on the precise interactions between the ribosome, the mRNA, and the nascent peptide in the ribosome exit tunnel (1, 45, 46). To determine the minimal elements necessary for efficient frameshifting, we conducted a series of mutational analyses focusing on both the nascent peptide sequence and the surrounding mRNA regions. Initially, we hypothesized that the sequences upstream of the frameshifting site, both within the nascent peptide and the mRNA, might play a role in modulating frameshifting efficiency. This hypothesis was based on the idea that these regions could influence the folding of the mRNA or the positioning of the ribosome in a manner that favors frameshifting. To test this, we introduced mutations into the upstream sequences of the nascent peptide and mRNA in our reporter constructs and measured the resulting frameshifting efficiency using the dual-luciferase assays described above (**Figure 1B**).

We deleted two uridine nucleotides close to the 5′ end of the minimal WNV frameshifting sequence, and inserted two uridines close to the slippery site. This deletion-insertion combination, which shifts the register of 121 nucleotides, changes the nascent peptide sequence with a minimal effect on the mRNA sequence (**Figure 2A** and **Figure S2**). These mutations did not lead to significant changes in frameshifting efficiency (**Figure 2B**), indicating that the nascent peptide translated prior to frameshifting does not play a critical role in the frameshifting process, despite its conservation upstream of the slippery site among WNV isolates (**Figure 2C**). Additionally, the complete deletion of the viral sequence upstream of the slippery site (5′ truncation of 123 nucleotides) also has little effect on the frameshifting efficiency. By contrast, deletion of sequences downstream of the frameshifting site 3′ of the predicted pseudoknot suppresses frameshifting substantially (**Figure 2D, E** and **Figure S3A**). These results show that the WNV -1 frameshifting site is dependent mainly on the slippery site and downstream sequences, which includes the RNA sequence thought to form a pseudoknot structure that induces ribosomal pausing.

**Figure 2.**
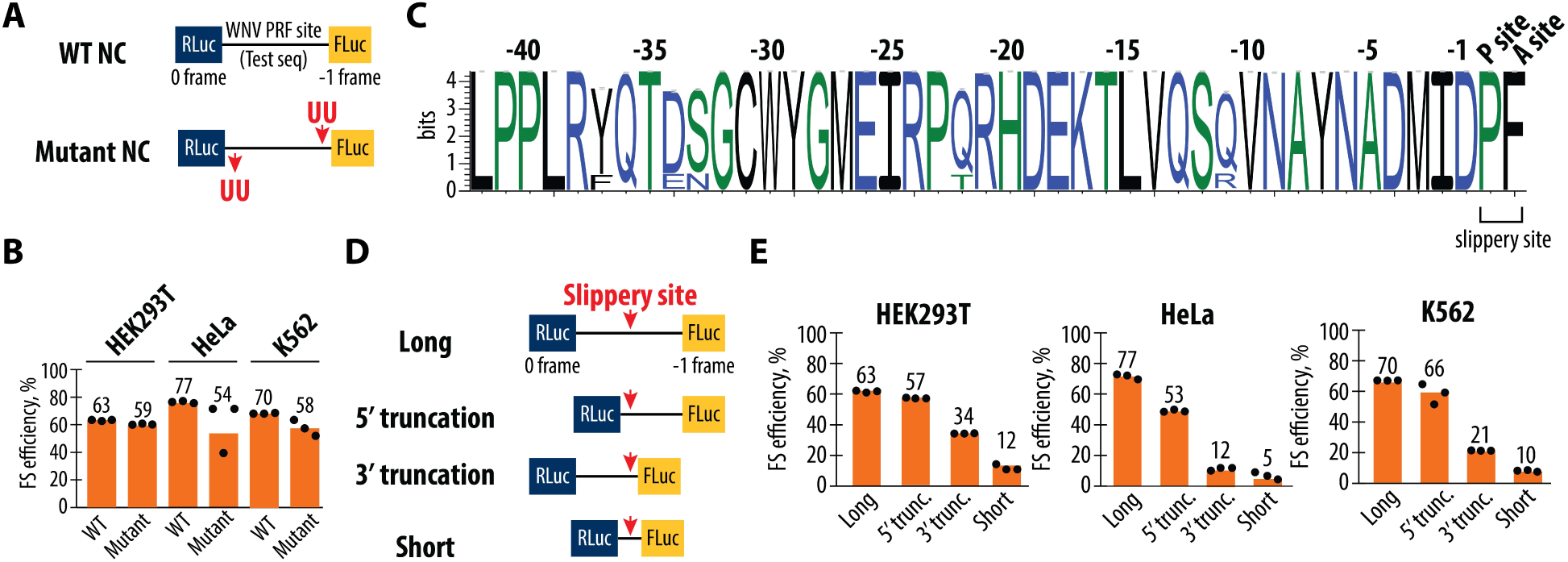
Importance of the nascent peptide and mRNA length for the ribosome frameshifting. (**A**) Schematic representation of the wild-type (WT) and mutant 2-nucleotide register shifted reporter constructs. In the WT construct, the firefly luciferase (FLuc) gene is positioned in the -1 reading frame relative to the Renilla luciferase (RLuc) gene, separated by the WNV PRF site. In the mutant register-shifted construct, two U residues deleted at the beginning of the testing sequence and inserted back upstream of the frameshifting site change the nascent peptide sequence while having a minimal effect on the mRNA sequence. Five additional nt changes that remove potential stop codons in the 2-nt register shifted sequence are shown in **Figure S2**. (**B**) Frameshifting efficiencies of the WT and mutant constructs measured in HEK293T, HeLa, and K562 cell lines. Frameshifting efficiency is shown as a percentage of the ratio between normalized FLuc/RLuc luminescence in the -1 frame to the sum of normalized Fluc/Rluc luminescence in the -1 and 0 frames. Bars represent the means of three independent experiments, with each dot corresponding to the individual experimental data points. (**C**) Sequence logo information content in bits of the 0-frame nascent peptide multiple sequence alignment immediately upstream from the WNV PRF site, for all complete WNV genomes deposited in the NCBI sequence database. The height of each position represents the relative information content of each position in bits and the x-axis displays the relative position of the amino acid in the multiple sequence alignment. (**D**) Schematic of truncation constructs used to assess the effect of sequences surrounding the slippery sequence site on frameshifting efficiency. The “Long” construct includes the full-length PRF sequence (258 nt), while the “5′ truncation,” (135 nt) “3′ truncation,” (204 nt) and “Short” (81 nt) constructs progressively reduce the sequence length around the slippery site. (**E**) Frameshifting efficiencies for the truncation constructs in HEK293T, HeLa, and K562 cells. Frameshifting efficiency is shown as a percentage of the ratio between normalized FLuc/RLuc luminescence in the -1 frame to the sum of normalized Fluc/Rluc luminescence in the -1 and 0 frames. Bars represent the means of three independent experiments, with each dot corresponding to the individual experimental data points.

Taken together, our results demonstrate that the sequences of the nascent peptide and mRNA upstream of the slippery sequence are not critical for frameshifting, while the downstream mRNA sequences play a vital role despite the high degree of the sequence conservation of the mRNA and polypeptides through the entire WNV -1 frameshifting site tested here (**Figure 2C** and **Figure S3B, C**). These findings suggest that the minimal requirements for the WNV frameshifting site include a functional slippery site followed by sequences capable of forming a stable RNA structure (i.e. a predicted pseudoknot), which together create the necessary conditions for efficient ribosomal frameshifting.

### The WNV Frameshifting Site Forms a Functionally Important and Sequence-Conserved Pseudoknot Structure

The WNV frameshifting site is predicted to form an H-type pseudoknot structure (25), as a key element in the frameshifting mechanism. Pseudoknots are thought to cause ribosomal pausing, which facilitates the slippage of tRNAs on the mRNA template, leading to frameshifting (47–49). However, in some viruses, pseudoknot formation is not required, and an RNA hairpin suffices to stimulate frameshifting (1, 46, 50). Furthermore, the WNV sequence downstream of the slippery site has been proposed to form multiple alternative RNA secondary structures that might contribute to the high frameshifting efficiency of the WNV PRF site (**Figure S4**) (27). However, sequence conservation and covariation analysis among flaviviruses show that mutations in the pseudoknot structure tend to be compensated by other mutations that maintain its integrity (**Figures S5** and **S6**). Additionally, a reanalysis of results of previously published structural probing data *in vitro* (25) and *in vivo* (51) also support the conclusion that the pseudoknot is the only likely mRNA secondary structure that can stimulate ribosome frameshifting (**Figure S7**). Nucleotides forming the stem of the pseudoknot showed reduced reactivity, while nucleotides in the bulge and loop regions displayed higher reactivity (**Figure S7A**). Additionally, nucleotides downstream from the predicted pseudoknot structure sequence proposed to form alternative structures (27) exhibited high reactivity, suggesting that these nucleotides are unlikely to form any stable structures. We notice an overall higher-than-expected reactivity of nucleotides involved in the formation of the predicted WNV pseudoknot, which may reflect both its intrinsic dynamic nature and the fact that the pseudoknot must undergo folding-unfolding cycle each time ribosome bypasses this mRNA region upon translation (**Figure S7**). Taken together, the sequence and structural probing analysis raise questions about what RNA structural elements are required for efficient WNV frameshifting.

To understand the role of the predicted pseudoknot in WNV -1 frameshifting, we performed a detailed mutational analysis aimed at disrupting the pseudoknot’s structure and assessed the impact on frameshifting efficiency. The pseudoknot structure is formed by base-pairing interactions between regions of the mRNA brought into close proximity by the folding of the RNA to form a long helix along with a pseudoknot RNA helix at the apex of the RNA stem-loop (46). To disrupt the pseudoknot, we introduced a series of mutations into the mRNA predicted to either weaken or completely abolish these base pairing interactions (**Figure 3A**). The frameshifting efficiency of these mutated constructs was then measured using the dual-luciferase assays developed in **Figure 1B**. Frameshifting efficiency was significantly reduced in all constructs where the pseudoknot’s stem regions including the pseudoknot base pairs were disrupted (constructs 2, 3, 5, and 6), consistent with the RNA structure playing an essential role in promoting efficient frameshifting (**Figure 3B**). The only mutations that were tolerated without affecting frameshifting efficiency were those in the internal loop region that links the primary helix to the predicted pseudoknot duplex (construct 4), where frameshifting efficiency remained comparable to the wild-type (WT) pseudoknot. These results support the model that the WNV frameshift-stimulating sequence forms a pseudoknot and that maintaining the integrity of the stems is critical for efficient frameshifting.

**Figure 3.**
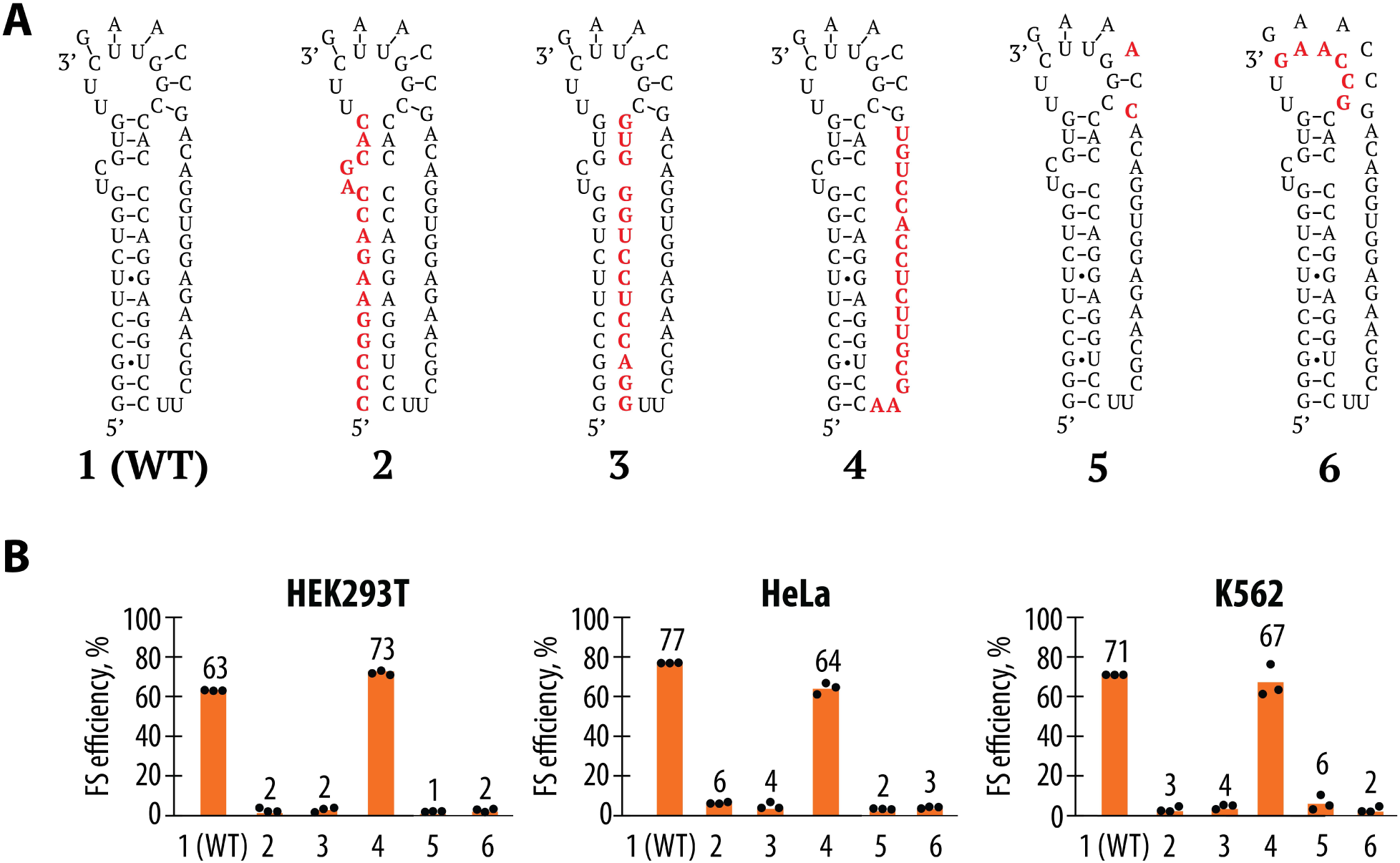
Mutational probing of the pseudoknot structure in WNV -1 frameshifting. (**A**) Schematic representation of the mutations designed to disrupt the predicted pseudoknot structure. Mutations are shown in red and were targeted to various regions of the pseudoknot, including the stem and loop regions. (**B**) Results of the frameshifting efficiency assays for the WT and mutant pseudoknot constructs in HEK293T, HeLa, and K562 cells. Frameshifting efficiency is shown as a percentage of the ratio between normalized FLuc/RLuc luminescence in the -1 frame to the sum of normalized Fluc/Rluc luminescence in the -1 and 0 frames. Bars represent the means of three independent experiments, with each dot corresponding to the individual experimental data points.

RNA secondary structures disrupted by mutations on one strand can often be rescued by mutations in the opposite strand that form compensatory base pairs. In an attempt to rescue the disrupted pseudoknot structure, we introduced compensatory mutations designed to restore base pairing in constructs 2, 3, 5, and 6. However, these compensatory mutations failed to restore frameshifting efficiency (**Figure S8**). This might suggest that the specific sequence of the WNV pseudoknot has been evolutionarily selected for its ability to form a single highly efficient frameshift-stimulating RNA structure. The extensive sets of mutations introduced here may be unable to fold correctly into a functional RNA pseudoknot structure due to alternative folds, for example. Notably, since these sequences are downstream of the slippery sequence site, the nascent peptide they encode cannot be the cause of the disrupted frameshifting we observe.

### Positioning of the Pseudoknot Downstream of the Slippery Site Required for Efficient WNV Frameshifting

In viral frameshifting sites, the RNA structural element downstream of the slippery site is thought to induce ribosome pausing to facilitate the frameshifting event (29, 30, 48, 49). In structures of the ribosome, the mRNA downstream of the P site threads through an mRNA entry tunnel over a span of about 16 nucleotides (where the +1 nt is the first nucleotide of P-site codon) (48, 52, 53). Thus, any RNA secondary structure 3′ of the first nucleotide of the P-site codon would be occluded by the ribosome if positioned closer than about 16 nucleotides (**Figure S9**). To investigate the importance of this spatial relationship in the context of WNV frameshifting, we engineered a series of reporter constructs with varying linker lengths between the slippery site and the pseudoknot. We tested four different linker lengths between 2 and 11 nts, with 5 nts corresponding to the WT sequence (**Figure 4A**). The WT linker length of 5 nt results in optimal frameshifting across all three cell lines (**Figure 4B**), suggesting that the distance between the slippery site and the pseudoknot in the WT sequence is evolutionarily optimized for efficient frameshifting. By contrast, a construct with a shorter linker (2 nt) showed significantly reduced frameshifting efficiency. Similarly, constructs with longer linkers (8 nt and 11 nt) also exhibited reduced frameshifting efficiency.

**Figure 4.**
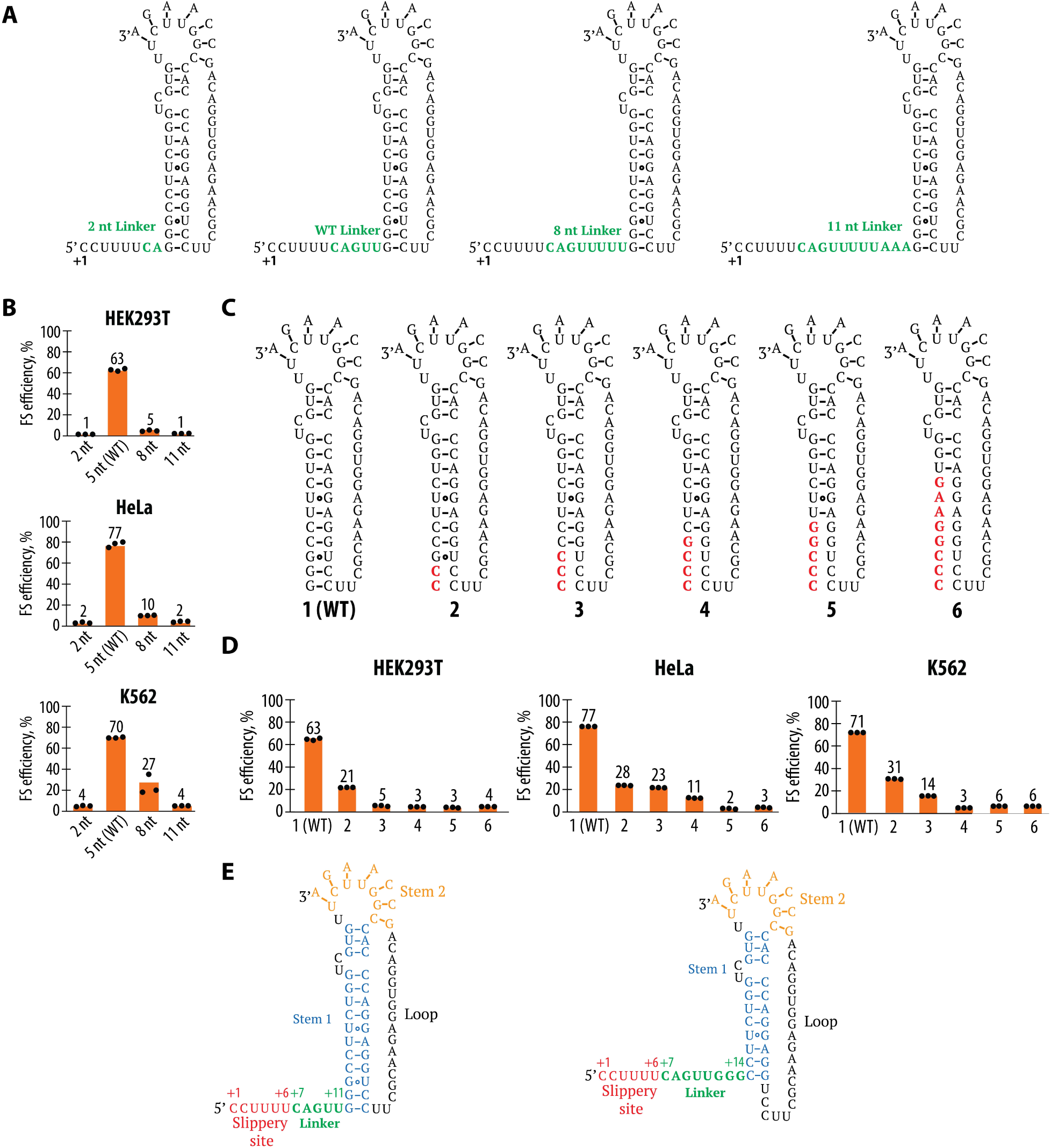
Analysis of Linker Length and Pseudoknot Structure of the WNV Frameshifting Site. (**A**) Schematic representation of the mutations altering the length of the linker region between the slippery site and the pseudoknot. (**B**) Frameshifting efficiency of the WNV sequence with different linker lengths in HEK293T, HeLa, and K562 cells. Frameshifting efficiency is shown as a percentage of the ratio between normalized FLuc/RLuc luminescence in the -1 frame to the sum of normalized Fluc/Rluc luminescence in the -1 and 0 frames. Bars represent the means of three independent experiments, with each dot corresponding to the individual experimental data points. (**C**) Representation of the WT pseudoknot structure and mutations that disrupt predicted base pairs at the base of the pseudoknot stem (mutations highlighted in red). Constructs are numbered 1 (i.e. WT) through 6. (**D**) Frameshifting efficiency of the mutant sequences with disrupted base pairs compared to WT, in HEK293T, HeLa, and K562 cells. Frameshifting efficiency is shown as a percentage of the ratio between normalized FLuc/RLuc luminescence in the -1 frame to the sum of normalized Fluc/Rluc luminescence in the -1 and 0 frames. Bars represent the means of three independent experiments, with each dot corresponding to the individual experimental data points. (**E**) Proposed structure of the WNV frameshifting site based on the reporter assays (right), compared to the previously predicted structure (left).

It is interesting to note that the optimal distance between the pseudoknot and slippery sequences corresponds to 5 nts, placing the predicted secondary structure for the pseudoknot at position +12 relative to the start of the P site. Even a linker of 8 nts (corresponding to the pseudoknot starting at position +15) might lead to steric clashes with the ribosome (**Figure S9**) (48, 52). We therefore introduced mutations that disrupt sets of base pairs at the base of the WNV pseudoknot to determine their importance for -1 frameshifting (**Figure 4C**). Our mutational analysis revealed that disruption of the first 4 to 8 base pairs in the pseudoknot stem resulted in a significant reduction in frameshifting efficiency (**Figure 4D**). Interestingly, when only the first 2-3 base pairs were mutated, frameshifting was still relatively robust, suggesting that the closing base pairs of the pseudoknot stem might actually serve as part of the linker, allowing for flexibility rather than providing stability to the pseudoknot structure. The observed effects of changing linker lengths and the base of the pseudoknot stem lead us to propose a new model of the WNV pseudoknot structure (**Figure 4E**). In the canonical model (left panel), the pseudoknot was predicted to be a fully formed stable structure with two intact stems. However, we suggest that the first 3 base pairs of Stem 1 do not contribute significantly to frameshifting, allowing for a more flexible region near the slippery site (right panel). Interestingly, this updated model for the pseudoknot structure would still place the pseudoknot closer to the P site than predicted from ribosome structures with the SARS-CoV-2 frameshifting site (+14 linker vs. +16 linker in **Figure S9**) (48).

### Modifying the Slippery Sequence of the WNV Frameshifting Site to U_UUU_UUU Promotes -2 Frameshifting

The slippery sequence is a key element of known -1 frameshifting sites, characterized by a sequence that allows the ribosome’s tRNAs to slip from one reading frame to another. The exact sequence of the slippery site has been shown to be critical in determining the frameshifting outcome in other contexts (36, 54). To explore the effects of different slippery sequences on WNV frameshifting, we modified the WT slippery sequence (C_CCU_UUU) to U_UUU_UUU, a sequence that naturally appears in some *Flaviviridae* viruses (**Figure S10**) (1). We introduced this modified sequence into our reporter constructs and measured frameshifting efficiency in HEK293T, HeLa, and K562 cells using the dual-luciferase assays. The U_UUU_UUU sequence maintained significant -1 frameshifting efficiency across all tested cell lines, although the efficiency was slightly reduced compared to the WT sequence (**Figure 5A, B**). Notably, the U_UUU_UUU sequence also induced substantial -2 frameshifting, a phenomenon not observed with the WT sequence (**Figure 5C, D**). The observation of -2 frameshifting suggests that the U_UUU_UUU sequence creates a different set of interactions between the tRNAs and the mRNA, allowing for an alternative frameshifting event. This finding indicates that the specific sequence of the slippery site plays a crucial role in determining the type of frameshifting that occurs, with certain sequences capable of promoting both -1 and -2 frameshifting.

**Figure 5.**
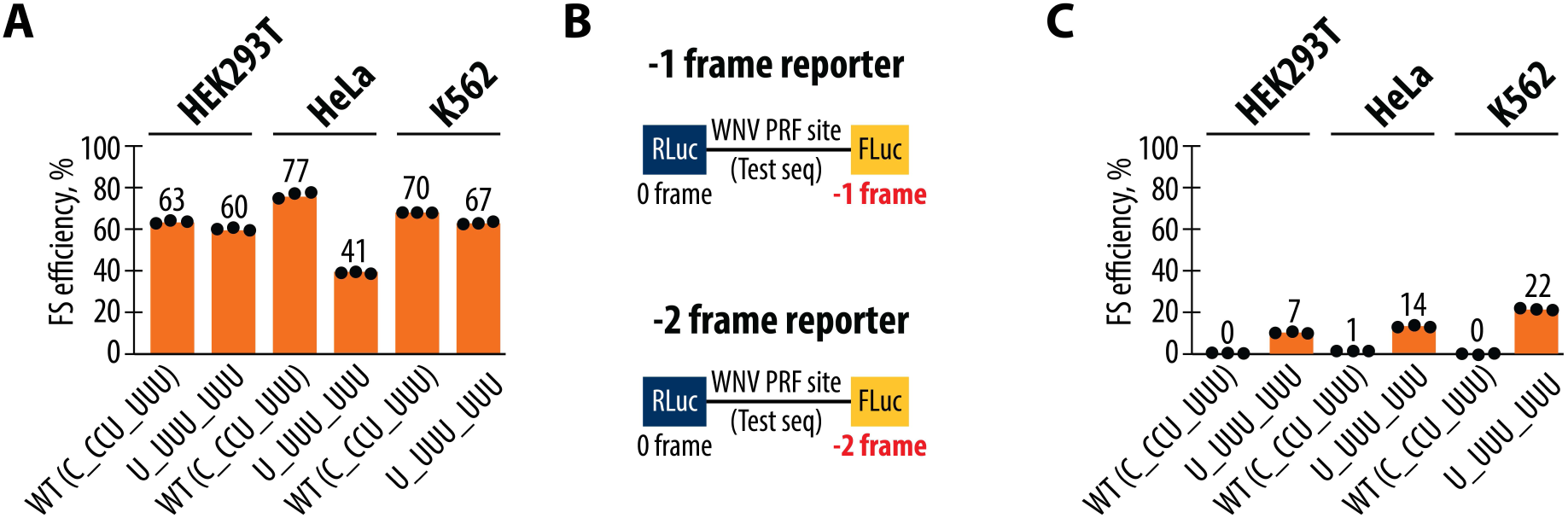
Effect of slippery site sequences on frameshifting efficiency and register in the West Nile Virus PRF site. (**A**) Frameshifting efficiencies of the WT (C_CCU_UUU) and mutant (U_UUU_UUU) frameshifting constructs in HEK293T, HeLa, and K562 cell lines. Frameshifting efficiency is shown as a percentage of the ratio between normalized FLuc/RLuc luminescence in the -1 frame to the sum of normalized Fluc/Rluc luminescence in the -1 and 0 frames. (**B**) Schematic representation of the dual-luciferase reporters designed to measure -1 and -2 ribosomal frameshifting. In the -1 frame reporter, FLuc is positioned in the -1 frame, whereas in the -2 frame reporter, FLuc is positioned in the -2 frame relative Rluc in the 0 frame. (**C**) Efficiency of -2 ribosomal frameshifting in HEK293T, HeLa, and K562 cells for the WT and mutant constructs. The WT sequence shows no detectable -2 frameshifting, while the mutant (U_UUU_UUU) construct induces low levels of -2 frameshifting, particularly in K562 cells. Frameshifting efficiency is shown as a percentage of the ratio between normalized FLuc/RLuc luminescence in the -2 frame to the sum of normalized Fluc/Rluc luminescence in the -2 and 0 frames. In panels (**A**) and (**C**), bars represent the means of three independent experiments, with each dot corresponding to the individual experimental data points.

### Role of the P-site and A-site slippery sequence base pairs in WNV Programmed Ribosome Frameshifting

A change to the slippery sequence that alters the P-site codon in both frames was sufficient to induce multiple frameshifting registers. This effect also raises questions about when frameshifting occurs, i.e. immediately after peptide bond formation, or after the mRNA and tRNAs are translocated by one codon by eEF2. We therefore conducted a more detailed mutational analysis focusing on the seven nucleotides directly involved in the frameshifting process, the codons in the ribosomal A and P sites and the first nucleotide of the slippery sequence which corresponds to the last nt of the E site in the 0 frame (**Figure 6**). We introduced specific mutations into the P-site and A-site codons (**Figure 6B** and **6D**, respectively) that would retain base pairing in the first, second, or third nucleotide of each codon prior to frameshifting, but that would alter their pairing after -1 frameshifting. Notably, these changes would also affect which tRNAs are used in translating the 0 frame, and the predicted number of mispaired codon-anticodon nucleotides after frameshifting occurs. Remarkably, disruption of tRNA-mRNA base pairs in the P site or in the first position of the A site after -1 frameshifting is not always deleterious to frameshifting efficiency, assuming that frameshifting occurs immediately after peptide bond formation but before translocation. For example, the second construct (**Figure 6B**) is predicted to fully pair in the 0 frame but results in mispairing at two positions in the P site codon after a -1 frameshift, yet still induces substantial frameshifting in the -1 frame (**Figure 6C**). Construct P4, which is designed to pair with tRNA^Pro^ in the P site mediated by the inosine post-transcriptional modification in the tRNA anticodon, would lead to a disrupted base pair in the first position of the A site codon (**Figure 6B**), yet retains almost WT levels of -1 frameshifting. By contrast, construct P5 differs from construct P4 in only the type of mispair in the first position of the A site codon in the -1 frame (**Figure 6B**), and results in very little -1 frameshifting (**Figure 6C**). Taken together, these results suggest that if frameshifting occurs before translocation, maintenance of P-site codon-anticodon interactions after -1 frameshifting is not necessary for frameshifting to occur, but the presence or the identity of mismatched bases in the first position of the A site in the -1 frame can dramatically affect whether frameshifting occurs (**Figure 6C**).

**Figure 6.**
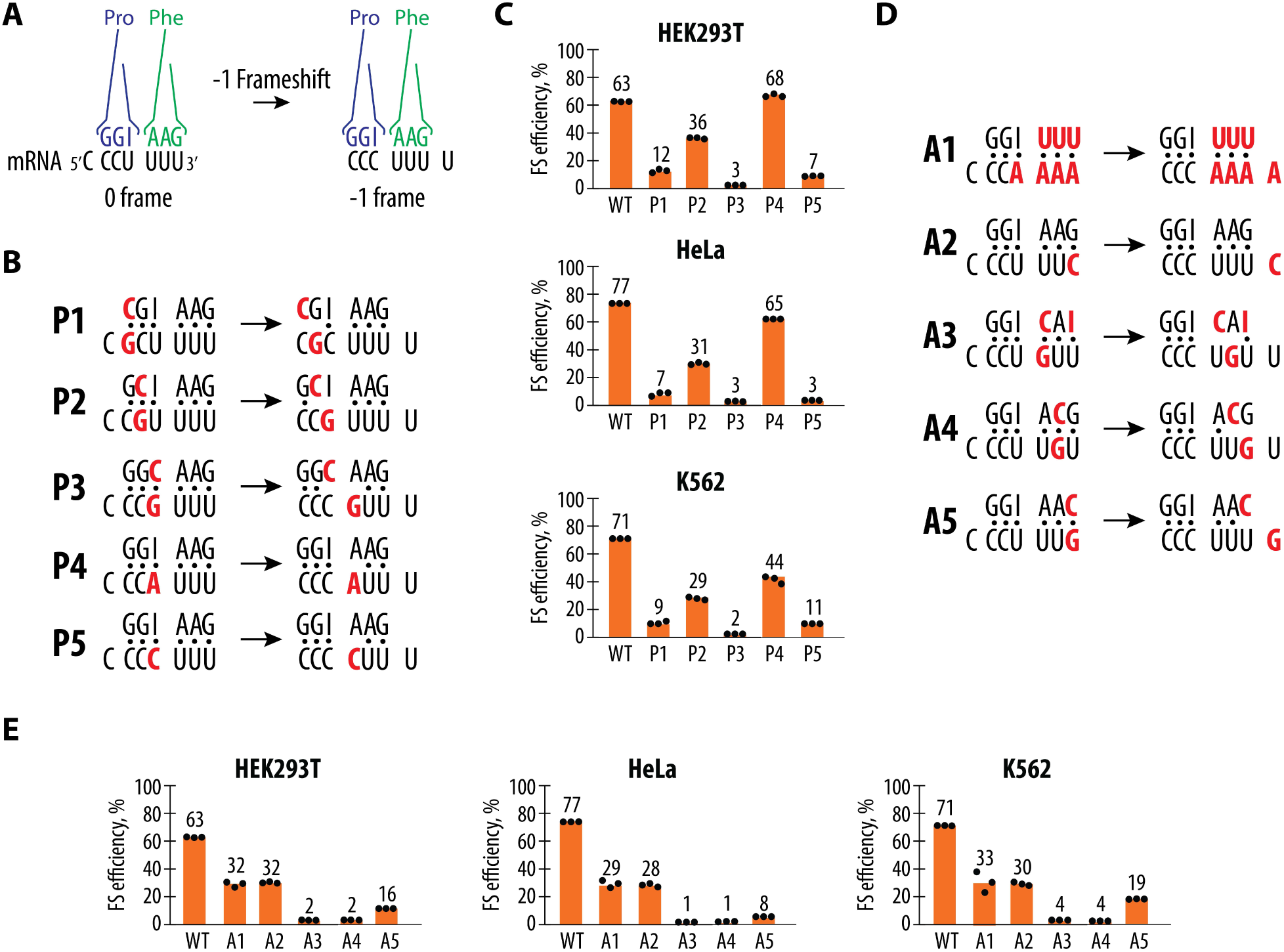
Mutational Analysis of Codon-Anticodon Interactions during WNV frameshifting. (**A**) Overview of codon-anticodon interactions before and after the -1 frameshift. P-site tRNA^Pro^(blue) and A-site tRNA^Phe^(green) in the example of the wild type WNV slippery site are shown interacting with their codons, shown in yellow. After frameshifting, the tRNA identities remain the same, but their positions on the mRNA shift. (**B**) Codon-anticodon interactions for constructs with mutations in the P-site codon before and after frameshifting. The mRNA mutations and corresponding changes in the anticodon sequences of tRNAs are shown in red. (**C**) Frameshifting efficiency for the P-site mutant constructs in HEK293T, HeLa, and K562 cells. Frameshifting efficiency is shown as a percentage of the ratio between normalized FLuc/RLuc luminescence in the -1 frame to the sum of normalized Fluc/Rluc luminescence in the -1 and 0 frames. (**D**) Codon-anticodon interactions for constructs with mutations in the A-site codon before and after frameshifting. The mRNA mutations and corresponding changes in the anticodon sequences of tRNAs are shown in red. (**E**) Frameshifting efficiency for the A-site mutant constructs in HEK293T, HeLa, and K562 cells. Frameshifting efficiency is shown as a percentage of the ratio between normalized FLuc/RLuc luminescence in the -1 frame to the sum of normalized Fluc/Rluc luminescence in the -1 and 0 frames. In panels (**C**) and (**E**) the bars represent the means of three independent experiments, with each dot corresponding to the individual experimental data points.

Previous structural and biochemical studies have shown that the ribosome can tolerate significant deviations in the geometry of codon-anticodon interactions in the first position of the P site (55). This could explain why mutations in the first nucleotide of the slippery site do not affect frameshifting efficiency, even though codon-anticodon interactions are compromised after frameshifting (**Figure S11**). A naturally occurring mutation (**U**_CCU_UUU) in the slippery sequence (**Figure S10**), according to our data (**Figure S11**), does not affect frameshifting efficiency as much as previously reported in the literature (25). This discrepancy may reflect differences in experimental conditions or the intrinsic flexibility of the WNV frameshifting mechanism, which appears capable of tolerating certain variations without losing functionality. However, the flexibility in the first position of the P site cannot explain the frameshifting efficiency of constructs that disrupt 2-3 base pairs in the P site, or the third position of the P site, after frameshifting (P1-P3 in **Figure 6B, C**). A more likely scenario is that frameshifting occurs during or after mRNA and tRNA translocation by eEF2. In this case, all the mispaired nts in constructs P1-P3 would reside in the E site, which does not require mRNA and tRNA base pairs to form (56–60).

Unlike the P site, the effects of mutations in the A site in the 0 frame on frameshifting efficiency appear to better correlate with codon-anticodon interactions after frameshifting (**Figure 6D, E**). Assuming frameshifting occurs after peptide bond formation but before translocation, in constructs retaining full codon-anticodon pairing (i.e., A1 and A2), -1 frameshifting efficiency is only partially reduced. By contrast, frameshifting efficiency is abolished when A-site codon mutations disrupt two codon-anticodon pairs in the A site after frameshifting (constructs A3-A4) and is severely reduced if the third codon position becomes unpaired after frameshifting (construct A5). Taken together, these results are more consistent with the model that frameshifting occurs after mRNA and tRNA translocation, in which the P site of the ribosome is much more selective to correct codon-anticodon interactions in the second and third codon positions in the -1 frame for efficient frameshifting (**Figure 6D, E**), with the first position also contributing (constructs P4 and P5 in **Figure 6B, C**).

## Discussion

Programmed ribosomal frameshifting, which changes the register of the mRNA open reading frame, is central to the life cycle of many RNA viruses. Here we probed the mechanism of PRF used by the West Nile Virus, one of the most efficient frameshifting events known in biology. Although PRF events require the positioning of two key elements in the mRNA, a slippery site spanning two adjacent codons and a downstream RNA structural element (61, 62), how these elements contribute to the highly efficient frameshifting of the WNV PRF site have not been elucidated. We show that successful frameshifting by WNV requires the precise coordination of three key RNA elements: the slippery site, a downstream pseudoknot structure, and the specific distance between these components. The high efficiency of the WNV frameshifting site also allowed us to probe in more depth variations in these three elements that can support efficient frameshifting, or that lead to multiple frameshifting registers (-1 and -2 frames).

One of the most crucial aspects of frameshifting is the requirement of an RNA structure downstream of the slippery site to induce the ribosome to frameshift. This RNA structural element likely acts as a temporary roadblock, by slowing down the ribosome (46, 48). The downstream RNA structure is generally predicted to be a pseudoknot, but RNA hairpins can be sufficient to induce frameshifting (1, 46, 50). The WNV PRF site has been proposed to form a pseudoknot in the downstream sequence (22), but alternative RNA structures have also been proposed based on single-molecule experiments (27). Using sequence covariation analysis, re-examining structural probing data (25) (51), and mutational analysis, we provide evidence that the pseudoknot is likely the major RNA structure in the WNV sequence that promotes efficient ribosome frameshifting. Disrupting the paired regions in the predicted pseudoknot, including the pairing to the loop, led to loss of frameshifting. By contrast, mutation of the long loop between the stem and pseudoknot pairs had little effect. These mutational data are only compatible with the pseudoknot structure, and not with other proposed RNA structural models.

The correct distance, or linker length, between the slippery site and downstream pseudoknot has also been shown to be essential for frameshifting efficiency in other systems such as in coronaviruses. A specific linker length may regulate a tension force between the codons in the P and A sites and the downstream RNA structure mediated by the ribosome mRNA entry channel, thereby inducing slippage of the mRNA during frameshifting (63, 64). Interestingly, previous structural models propose that the distance from the P-site codon +1 position to the RNA structure spans 16 nts (48). In structures of the SARS-CoV-2 PRF site on the rabbit ribosome, the downstream pseudoknot could only be seen when located at +17 in the mRNA. Shorter distances resulted in structures that lacked defined density for the pseudoknot, either due to increased RNA dynamics or due to RNA unfolding by the ribosome (48). Here, we show that the WNV pseudoknot is positioned closer to the P site, potentially as close as the +12 nt. Notably, however, the base pairs at the base of the pseudoknot stem may only form transiently, if at all, in the WNV frameshifting mechanism. With these transient base pairs disrupted, the pseudoknot would be positioned at nucleotide +15 rather than +17 seen with the SARS-CoV-2 PRF site (48), which may be important for inducing tension between the ribosome and mRNA required for efficient frameshifting.

In viral PRF, a slippery site induces a -1 frameshift, allowing the ribosome to “slip” back by one nucleotide after reading the codons in the slippery site. However, the mechanism by which the slippery site sequence contributes to frameshifting has not been delineated. We were able to use the high efficiency of WNV frameshifting to probe the mechanism in more depth. We explored mutations at all seven positions of the slippery site, which help distinguish between three potential models for frameshifting that could occur during different stages of the translation elongation cycle (**Figure 7**). In one scenario, frameshifting could occur immediately after peptide bond formation, but before translocation of the mRNA by one codon by eEF2 (**Figure 7B**). Our data reveal three examples in which such a mechanism would lead to 1-2 mispaired bases in the P site and A site codons (for instance, constructs P2, P4, and A5 in **Figure 6**). Given the stringent requirements for base pairing between mRNA and tRNAs (55, 65, 66) these data argue against frameshifting before translocation. Alternatively, if frameshifting occurs during or after translocation, then in the case of construct P2 the mispairing that occurs in the -1 frame would occur in the ribosomal E site (**Figure 7B**), which does not require codon-anticodon base pairing (55–60). For constructs P4 and A5, one mispair would occur in the P site rather than in the A site, while possibly maintaining some base pairs in the E site. Furthermore, if frameshifting occurs during or after translocation, the P4 and A5 constructs would require translation elongation to proceed once with a mispaired tRNA (once in the P site), rather than twice (once in the A site, and a second time in the P site in the next round of translation elongation). Our data so far cannot rule out mechanisms of frameshifting during or after translocation. In either case, the final state of the ribosome would involve one tRNA in the E site, and the second in the P site in the -1 frame. The mutations we have identified that permit substantial frameshifting in the WNV PRF site should enable future experiments to unravel the molecular mechanism underlying the frameshifting event.

**Figure 7.**
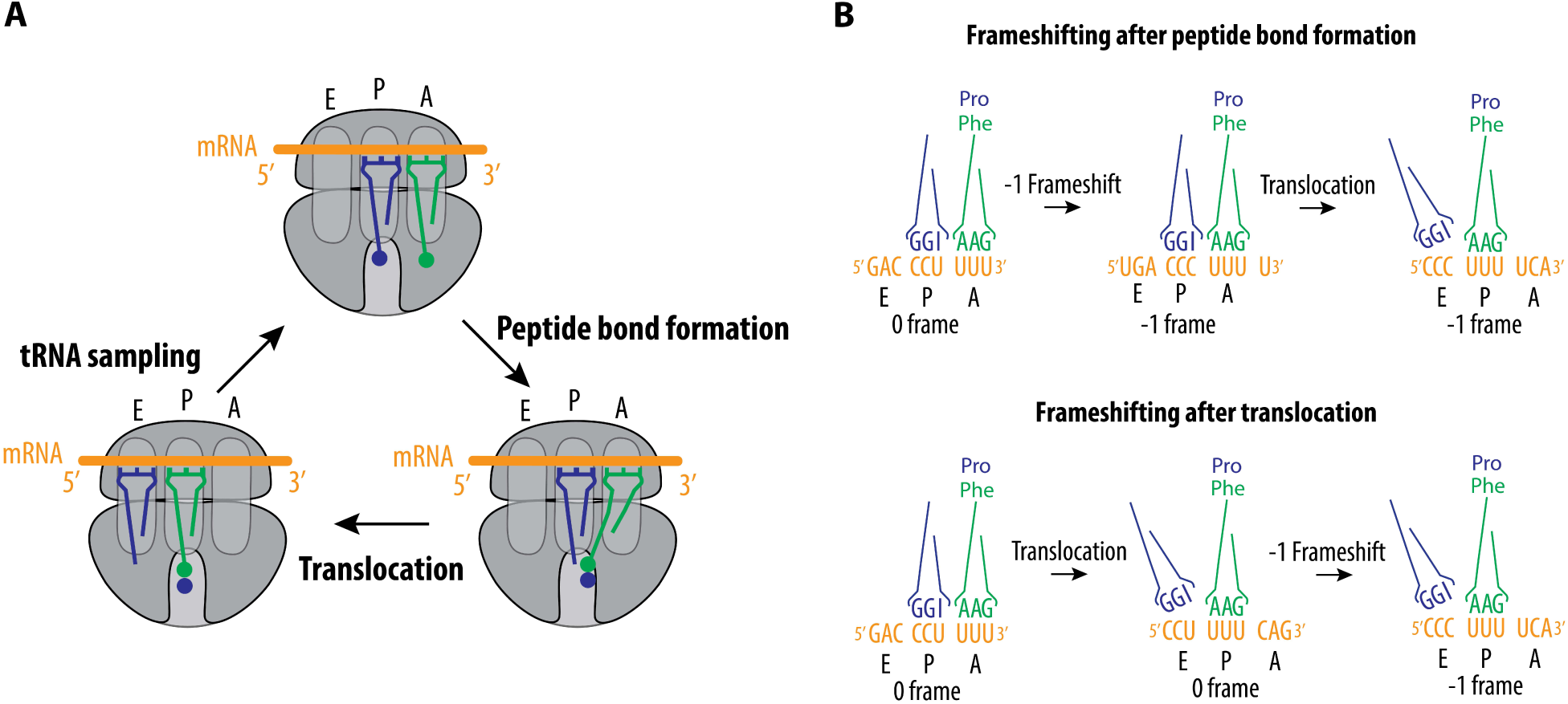
Model of WNV -1 frameshifting. (**A**) The regular elongation cycle consists of tRNA sampling, peptide bond formation, and translocation, with ribosomes spontaneously changing reading frames at a very low rate (∼0.01%) (67–69). (**B**) Possible models of frameshifting induced by the WNV PRF site: (1) Frameshifting may happen after peptide bond formation but before the ribosome undergoes translocation. (2) Frameshifting may alternatively occur after translocation and E-site tRNA dissociation, when only one tRNA occupies the P-site. This would establish the -1 frame for the next mRNA decoding event in the A site. Not shown, frameshifting could also take place simultaneously with the translocation event, resulting in the ribosome shifting by one nucleotide to the -1 frame, with the tRNAs positioned in the E site and P site.

In addition to insights into the step of elongation during which frameshifting occurs, our analysis of slippery site sequences revealed that these sites can promote frameshifting into multiple frames. The ability of the U_UUU_UUU sequence to promote -2 frameshifting highlights the malleability of the WNV frameshifting mechanism. Depending on the sequence context, the frameshifting mechanism can lead to different outcomes, with the potential for producing alternative proteins that may have distinct functions in the viral life cycle. This aligns with previous studies, which reported that different slippery site sequences can influence the extent and direction of frameshift events (70). The versatility of the slippery site in promoting both -1 and -2 frameshifting could also have important implications for some *Flaviviridae* strains in their ability to produce different proteins under various conditions, potentially contributing to its adaptability and virulence. Taken together, the level of frameshifting we observe across a wide variety of codon pairs suggests that frameshifting events may occur more frequently than previously appreciated. Searches for potential frameshifting sites may need to be expanded to account for this flexibility in frameshifting. Our results could also be used to define potential frameshifting sites in cellular genomes that may have previously been overlooked.

## Material and Methods

### Cell lines and culture conditions

HEK293T and HeLa cells were grown in DMEM media (Gibco) supplemented with 10% FBS (Gibco), penicillin (100 µg/ mL) (Gibco), streptomycin (100 µg/mL) (Gibco), and 2 mM GlutaMax (Gibco). K562 cells were grown in RPMI 1640 (Gibco) supplemented with 10% FBS (Gibco), penicillin (100 µg/ mL) (Gibco), and streptomycin (100 µg/mL) (Gibco). Cells were grown at 37 °C in 5% carbon dioxide and 100% humidity.

### Cloning

For PCR amplification, we used the Q5 High-Fidelity DNA Polymerase (NEB), and for mutagenesis, the Q5 Site-Directed Mutagenesis Kit (NEB). The Gibson assembly was performed using the NEBuilder HiFi DNA Assembly Master Mix (NEB). All PCR reactions, Q5 mutagenesis, Gibson assembly, and restriction digest reactions were performed according to the respective manufacturer’s instructions. All plasmids were verified through full plasmid sequencing (Plasmidsaurus). Primers were ordered from Integrated DNA Technologies (IDT), and synthetic DNA blocks were obtained from Twist Bioscience.

To generate the constructs described in **Figure 1B** (WT, IFC, SSM, 5’ STOP), the pSGDLuc v3.0 plasmid (Addgene, 119760) was digested with PspXI and BglII (both from NEB) to generate the vector backbone for cloning. The inserts were assembled using synthetic DNA blocks (Twist Bioscience) via Gibson assembly, with sequences provided in **Supplementary Table 1**. To generate the 3’ STOP construct described in **Figure 1B**, we used the WT construct generated in the previous step, followed by PCR amplification using primers listed in **Supplementary Table 2**. Gibson assembly was performed using an oligonucleotide as a fusing linker, with the sequence also listed in **Supplementary Table 2**. Constructs depicted in **Figures 3A, 4A, 4C, 5A, 5B, 6B, 6D, S8A**, and **S10A** were generated using the WT plasmid from **Figure 1B** as the template, and mutagenesis was performed with the Q5 Site-Directed Mutagenesis Kit according to the manufacturer’s instructions. Primer sequences can be found in **Supplementary Table 2**. To generate the constructs described in **Figure 1D**, the vector was generated by PCR using the primers from **Supplementary Table 2** with the previously described EMCV IRES-3x FLAG-GS-linker-NanoLuciferase plasmid (71, 72) as the template. The insert was amplified using primers from Supplementary Table 2 and WT/SSM plasmids (described in **Figure 1B**) as templates. DpnI (NEB) digestion was performed on PCR products before proceeding with Gibson assembly.

To generate the constructs described in **Figure 2A**, the pSGDLuc v3.0 plasmid (Addgene, 119760) was digested with ApaI (NEB) and then assembled with synthetic DNA blocks (sequences provided in Supplementary Table 1) via Gibson assembly. To generate the 5’ and 3’ truncation constructs described in **Figure 2D**, PCR amplification was performed on WT and IFC plasmids from **Figure 1B** using the primers listed in **Supplementary Table 2**, followed by ligation with the Q5 Site-Directed Mutagenesis Kit. To generate the short constructs depicted in **Figure 2D**, pSGDLuc v3.0 plasmid (Addgene, 119760) was digested with PspXI and BglII (both from NEB) to prepare the vector backbone. Inserts were generated using a 3-oligonucleotide PCR reaction, and the oligonucleotide sequences can be found in **Supplementary Table 2**. Gibson assembly was subsequently performed.

To generate the -2 reporters described in **Figure 5B**, the pSGDLuc v3.0 plasmid (Addgene, 119760) was digested with ApaI (NEB) to generate the vector. Gibson assembly was performed with synthetic DNA blocks, and their sequences are listed in **Supplementary Table 1**. The resulting plasmids contained the WT slippery sequences. To introduce mutations in the slippery site, we employed the Q5 Site-Directed Mutagenesis Kit with oligonucleotide sequences listed in **Supplementary Table 2**. The WT and IFC constructs in **Figure 1B** were also mutagenized in a similar fashion to generate U_UUU_UUU -1 frame reporters.

### Transfections and Dual-Luciferase assay

Cells were seeded in a 96-well format at a density of ∼10^4^ cells per well and transfected 24 h post-seeding with 100 ng reporter plasmid using Lipofectamine 3000 with P3000 reagent (Thermo Fisher Scientific), for HEK293T and HeLa cells, or Lipofectamine LTX with Plus reagent (Thermo Fisher Scientific) for K562 cells, according to the manufacturer’s instructions. Twenty-four hours after transfection, media was removed, cells were washed with DPBS (Gibco) and were lysed in 50 μl 1X passive lysis buffer (Promega). Cell extracts were frozen and thawed, and luciferase activity was measured using the Dual-Luciferase Reporter Assay System (Promega) by mixing 10 μl of cell extract with 50 μl of each reagent according to the manufacturer’s instructions. Background level readings (no transfection sample) were subtracted from all F-Luc and R-Luc readings. Frameshifting efficiencies were calculated according to the formula:

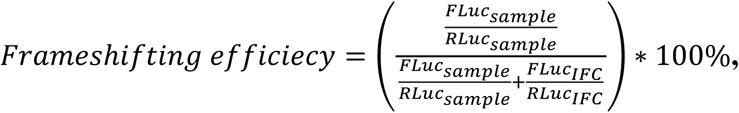

Where the sum in the denominator accounts for all translation events in both -1 and 0 frames.

### Western blot analysis

Samples were boiled in Bolt LDS Sample loading buffer (Thermo Fisher Scientific) containing Bolt/NuPAGE reduction buffer (Thermo Fisher Scientific) at 70 °C for 10 min. Samples were resolved on 4%–12% Bolt Bis-Tris Plus protein gels (Thermo Fisher Scientific) using 1× Bolt MES SDS running buffer (Thermo Fisher Scientific) containing 1× NuPAGE Antioxidant (Thermo Fisher Scientific), according to the manufacturer’s instructions. Gels were transferred to nitrocellulose membranes using a Power Blot system (Thermo Fisher Scientific) using medium-range manufacturer parameters. Membranes were blocked with 5% nonfat dry milk (BioWorld) in PBST (10 mM Tris-HCl, pH 8 [Invitrogen], 1 mM EDTA [Invitrogen], 0.1% Triton X-100 [Sigma], 150 mM sodium chloride [Sigma]) for 1 h at room temperature (RT) with gentle rocking. Blots were washed in PBST three times and then incubated with the indicated primary antibodies overnight at 4 °C with gentle rocking. Blots were washed with PBST three to four times over 45–60 min at RT with gentle rocking, then incubated with secondary antibodies diluted in 5% milk in PBST for 1 h with gentle rocking at RT. Membranes were washed again with PBST, three to four times over 45–60 min at RT, developed using SuperSignal West Pico Plus ECL substrate (Thermo Fisher Scientific) and SuperSignal West Femto Maximum Sensitivity substrate (Thermo Fisher Scientific) if needed and imaged on an iBright CL1000 (Thermo Fisher Scientific) system. Results shown are representative of at least two independent experiments. Frameshifting efficiency was quantified using ImageJ (73).

### Antibodies

The following antibodies were used in this study. Antibodies against the HA-tag (3724S) were from Cell Signaling Technology. Antibodies against RPS19 (A304-002A) were purchased from Bethyl Laboratories Inc. Anti-mouse IgG-HRP (sc-525409) was from Santa Cruz Biotechnology. Anti-FLAG-tag (F1804-50UG) was from Sigma. Anti-rabbit ECL IgG-HRP (NA934V) was from Thermo Fisher Scientific.

### In vitro transcription reactions

*In vitro* transcription reactions were performed using PCR products generated with primers encoding a flanking T7 RNA polymerase promoter:

(5’-TGGAGCTTAATACGACTCACTATAG-3’)

and a poly(A) tail:

(5’-T_39_CGTTGGGAGCTCTCCGGATCC-3’). Reactions were set up, as previously described (44), with 20 mM Tris-HCl, pH 7.5, 35 mM MgCl_2_, 2 mM spermidine, 10 mM DTT, 1 u/mL pyrophosphatase (Sigma), 7.5 mM of each NTP, 0.2 u/mL RiboLock RNase Inhibitor (Thermo Fisher Scientific), 0.1 mg/mL T7 RNA polymerase, and 40 ng/µL PCR-generated DNA. After 3 h incubation at 37 °C, 0.1 u/µL DNase I (Promega) was added to the reactions, which were incubated at 37 °C for 30 min to remove the template DNA. RNA was precipitated for 2–3 h at −20 °C after adding 0.5× volume of 7.5 M LiCl/ 50 mM EDTA, and the resulting pellet was washed with cold 70% ethanol and dissolved with RNase-free water. The mRNA was further purified by using a Zymo RNA Clean and Concentrator Kit (Zymo Research) before use in *in vitro* translation reactions.

### In vitro translation reactions

Translation reactions were set up using a high-efficiency human *in vitro* translation extract according to a previously published procedure (44). Briefly, for a 10 µL reaction, 5 µL of cell extract was used in a buffer containing final concentrations of 52 mM HEPES, pH 7.4 (Takara), 35 mM KGlu (Sigma), 1.75 mM Mg(OAc)_2_ (Invitrogen), 0.55 mM spermidine (Sigma), 1.5% Glycerol (Thermo Fisher Scientific), 0.7 mM putrescine (Sigma), 5 mM DTT (Thermo Fisher Scientific), 1.25 mM ATP (Thermo Fisher Scientific), 0.12 mM GTP (Thermo Fisher Scientific), 100 µM L-Arg; 67 µM each of L-Gln, L-Ile, L-Leu, L-Lys, L-Thr, L-Val; 33 µM each of L-Ala, L-Asp, L-Asn, L-Glu, Gly, L-His, L-Phe, L-Pro, L-Ser, L-Tyr; 17 µM each of L-Cys, L-Met; 8 µM L-Trp, 20 mM creatine phosphate (Roche), 60 µg/mL creatine kinase (Roche), 4.65 µg/mL myokinase (Sigma), 0.48 µg/mL nucleosidediphosphate kinase (Sigma), 0.3 u/mL inorganic pyrophosphatase (Thermo Fisher Scientific), 100 µg/mL total calf tRNA (Sigma), 0.8 u/µL RiboLock RNase Inhibitor (Thermo Fisher Scientific), and 1000 ng mRNA. Translation reactions were incubated for 60 min at 32 °C. A 1 µl aliquot was used for western blot analysis.

### Bioinformatic searches for flavivirus frameshifting elements

Bioinformatic searches for flavivirus frameshifting elements used the bioinformatic software suite Infernal v1.1.2 (74). First, datasets containing all *Flaviviridae* RefSeq genomes, all *Flaviviridae* sequences, all complete West Nile Virus genomes, and all West Nile Virus genomes irrespective of completeness were downloaded from the NCBI (retrieval date 4/1/2024). These were then searched using the flavivirus frameshifting element consensus model from Rfam 14.10 (RF01768) (75). Identified sequences were then filtered to ensure they were present on the positive sense strand, were not truncated, and were required to have an E-value less than 0.01. These sequences were then extracted using Easel (76) and realigned to the RF01768 model using Infernal’s cmalign function.

### In silico *Flaviviridae* frameshifting element covariation analysis

Covariation analysis was performed to assess potential supported alternative RNA folds for the *Flaviviridae* frameshifting element with surrounding sequence. The sequences identified from the search across all *Flaviviridae* described above were extracted including 300 nucleotides upstream and downstream of the frameshifting element. These sequences were then realigned to an extended version of the RF01768 model using Infernal. Additional covariation-supported RNA folds were then searched for using the CaCoFold function of R-Scape v2.0.4.a (77, 78). No alternative statistically supported folds were identified.

### West Nile Virus polyprotein translation

To generate a local alignment of the West Nile Virus polyprotein around the frameshifting element, 126 nucleotides upstream and downstream were extracted and translated into amino acids using the Biopython Bio.Seq package (https://biopython.org/docs/dev/api/Bio.Seq.html) using the human codon table.

### WebLogo generation

Sequence WebLogos were generated using the command line executable version of WorkLogo3 (79). Protein Logos were generated considering an alphabet of 21 letters, the 20 canonical amino acids and a stop codon character, and rendered using the standard hydrophobicity color scheme including red for stop codons. Nucleic acids were generated using a 4-letter alphabet and colored using the default nucleotide color scheme.

### West Nile Virus frameshifting element tree construction

Sequences for tree generation were taken from the *Flaviviridae* RefSeq search above and clustered at a 0.95 identity level using MMseqs2 (80). The frameshifting element from the SARS-CoV-2 RefSeq genome (MN908947.3) was included as an outgroup. These sequences were then aligned with MAFFT (81) 7.490 using the –auto flag. The phylogenetic tree was calculated using two methods to ensure topological consistency, using a maximum-likelihood approach in MEGAX (82) and a Bayesian approach using MrBayes 3.2.7a (83). The tree was then rendered using the ITOL webserver (https://itol.embl.de/) (84).

### West Nile Virus *in vivo* chemical probing data reprocessing

*In vivo* NAI chemical probing reactivities of the West Nile Virus genome from infected Vero cells were downloaded from the GEO Accession Database (GSE228446)(51). The reactivity profiles of the two replicates were averaged and plotted onto three pseudoknot containing regions of the West Nile Virus genome, the frameshifting elements and two dumbbell structures, using VARNA (85).

### Rendering of cryo-EM structure of SARS-CoV-2 PRF site on the rabbit ribosome

Graphics for the cryo-EM structure from PDB entry 7o7z (48) were made using PyMOL (The PyMOL Molecular Graphics System, Version 2.5.8 Schrödinger, LLC).

## Acknowledgments

We thank the members of the J.H.D.C. laboratory for the helpful discussion and Dr. Yekaterina Shulgina for the critical reading of the manuscript.

