## Supplementary Information for "RNA elements required for the high efficiency of West Nile Virus-induced ribosomal frameshifting"

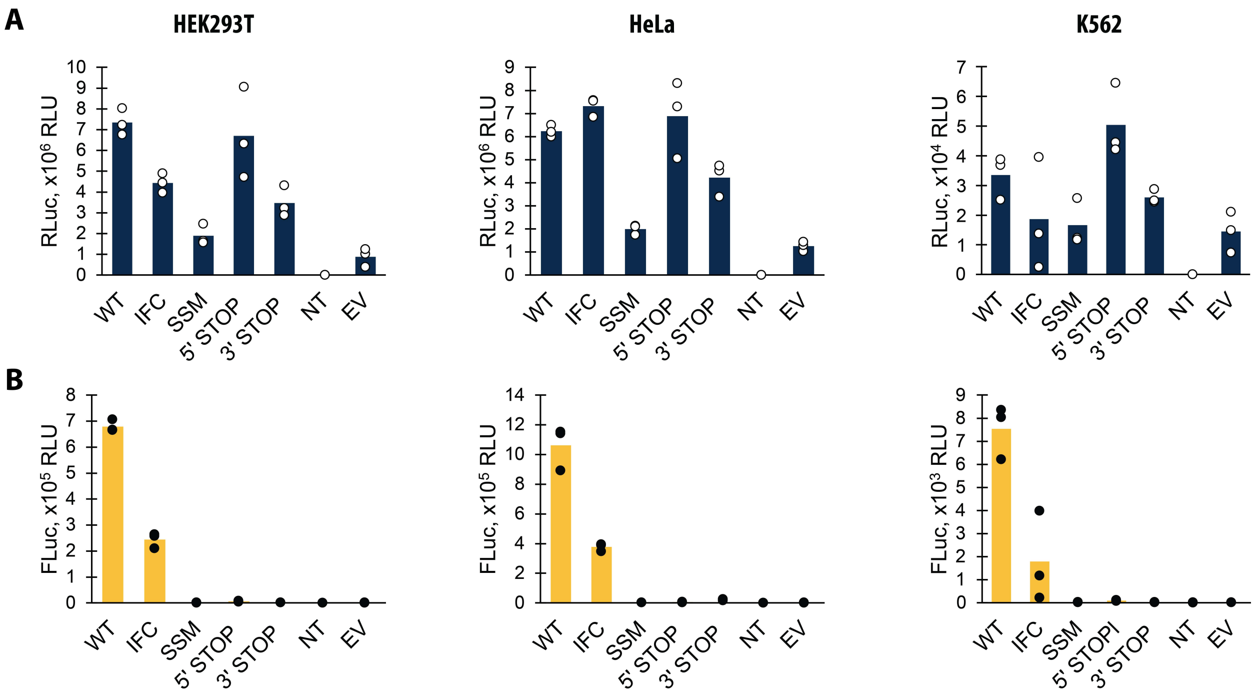


**Figure S1. Measurement of Renilla and Firefly luciferase activities in frameshifting reporter assays across different cell lines.**

(**A**) Renilla luciferase (RLuc) activities of the wild-type (WT) and mutant reporter constructs measured in HEK293T, HeLa, and K562 cell lines.

(**B**) Firefly luciferase (FLuc) activities measured from the same reporter constructs across the three cell lines. The WT and IFC constructs show high FLuc activity, with the IFC demonstrating efficient translation in the 0 frame. In contrast, the SSM, 5′ STOP, 3′ STOP, and EV constructs show minimal FLuc activity, consistent with impaired translation due to mutations or stop codon introduction at the frameshifting site.

Bars represent the means of three independent experiments, with each dot corresponding to the individual experimental data points.

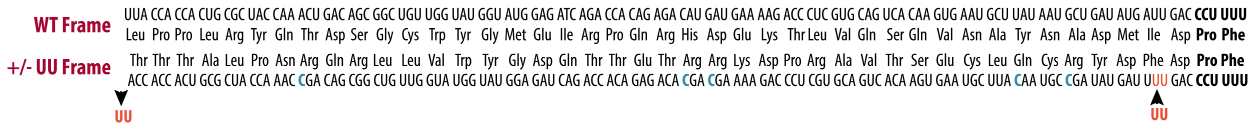


**Figure S2. Detailed representation of the nucleotide and amino acid sequences of the wild-type (WT) and mutant UU frameshifting constructs.**

This figure shows the nucleotide and corresponding amino acid sequences of the wild-type (WT) and mutant frameshifting constructs. In the mutant construct, two nucleotides (UU) were deleted near the 5′ end of the sequence and reinserted closer to the slippery site (marked with arrows). The slippery site is shown in bold. Five nucleotide changes made to remove stop codons that appeared after the UU deletion are shown in blue and bolded.

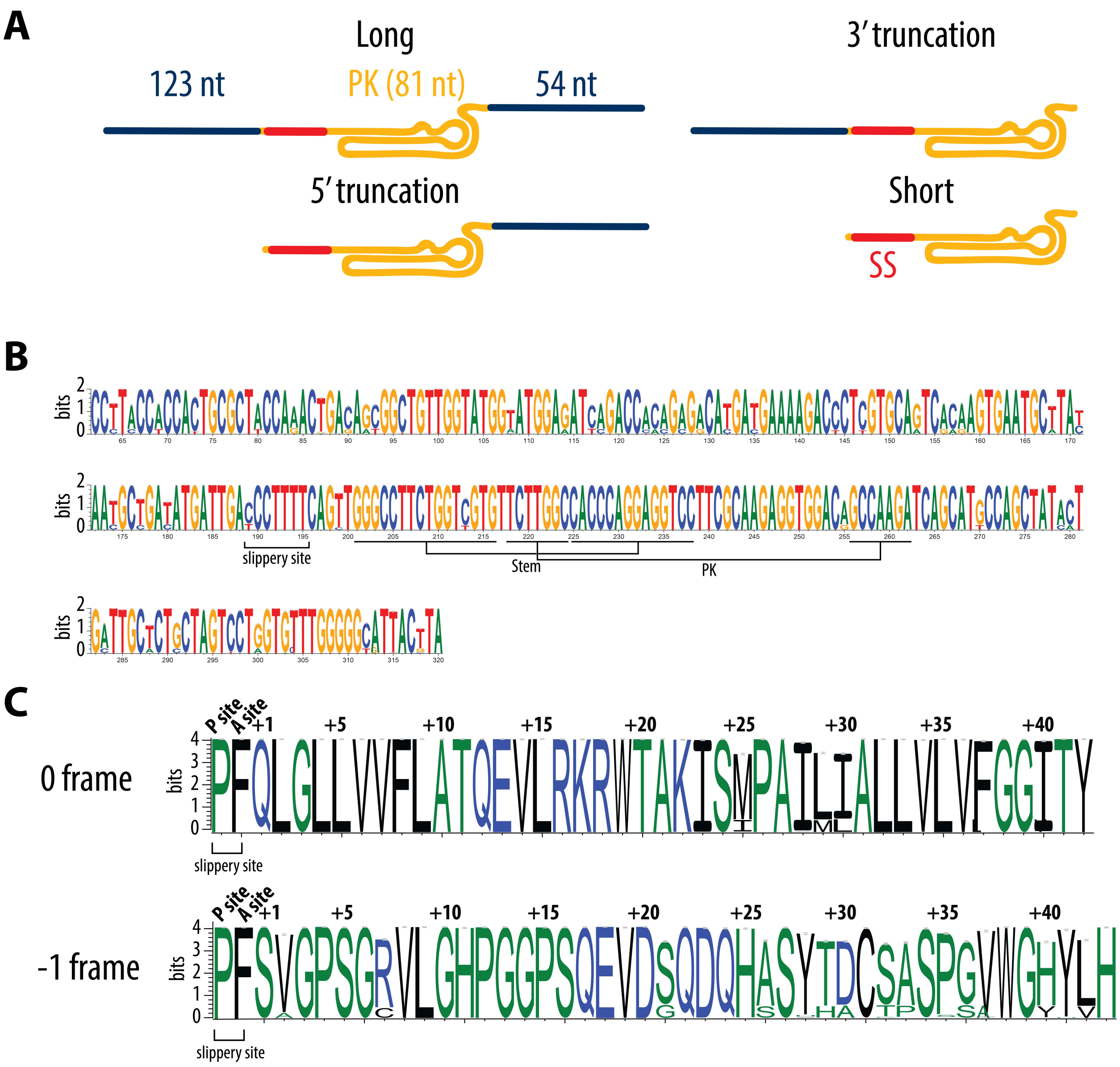


**Figure S3. Detailed schematic of truncation constructs and sequence conservation at the WNV PRF site.**

(**A**) Schematic representation of the truncation constructs related to **Figure 2D**. The blue regions indicate the flanking sequences with numbers of nucleotides, the golden regions represent the pseudoknot (PK) structure, and the red marks show the slippery site (SS). The "Long" construct includes the full-length flanking sequences and pseudoknot. The "5′ truncation" and "3′ truncation" constructs progressively reduce the flanking sequences, while the "Short" construct contains minimal flanking sequences with the pseudoknot and slippery site intact.

(**B**) Information content in bits of nucleotides upstream and downstream from the WNV programmed -1 ribosomal frameshifting (PRF) site were calculated using WebLogo3 for all complete WNV genomes deposited in the NCBI database. The height of each position represents the relative information content of each position in bits and the x-axis displays the relative position of the nucleotide in the multiple sequence alignment. The slippery site and nucleotides involved in pseudoknot formation are indicated. The sequence logo illustrates the degree of conservation for each nucleotide position, highlighting key elements necessary for frameshifting and predicted pseudoknot stability.

(**C**) Information content in bits of the translated polypeptides downstream from the frameshifting site in both the 0 frame and -1 frame calculated using WebLogo3 for all complete WNV genomes deposited on NCBI. The height of each position represents the relative information content of each position in bits and the x-axis displays the relative position of the amino acid in the multiple sequence alignment. The slippery site is indicated, and the sequence logo shows the conservation level of each amino acid in the two reading frames, providing insights into the potential variability of translation products depending on frameshifting events. Codon 44 is a stop codon.

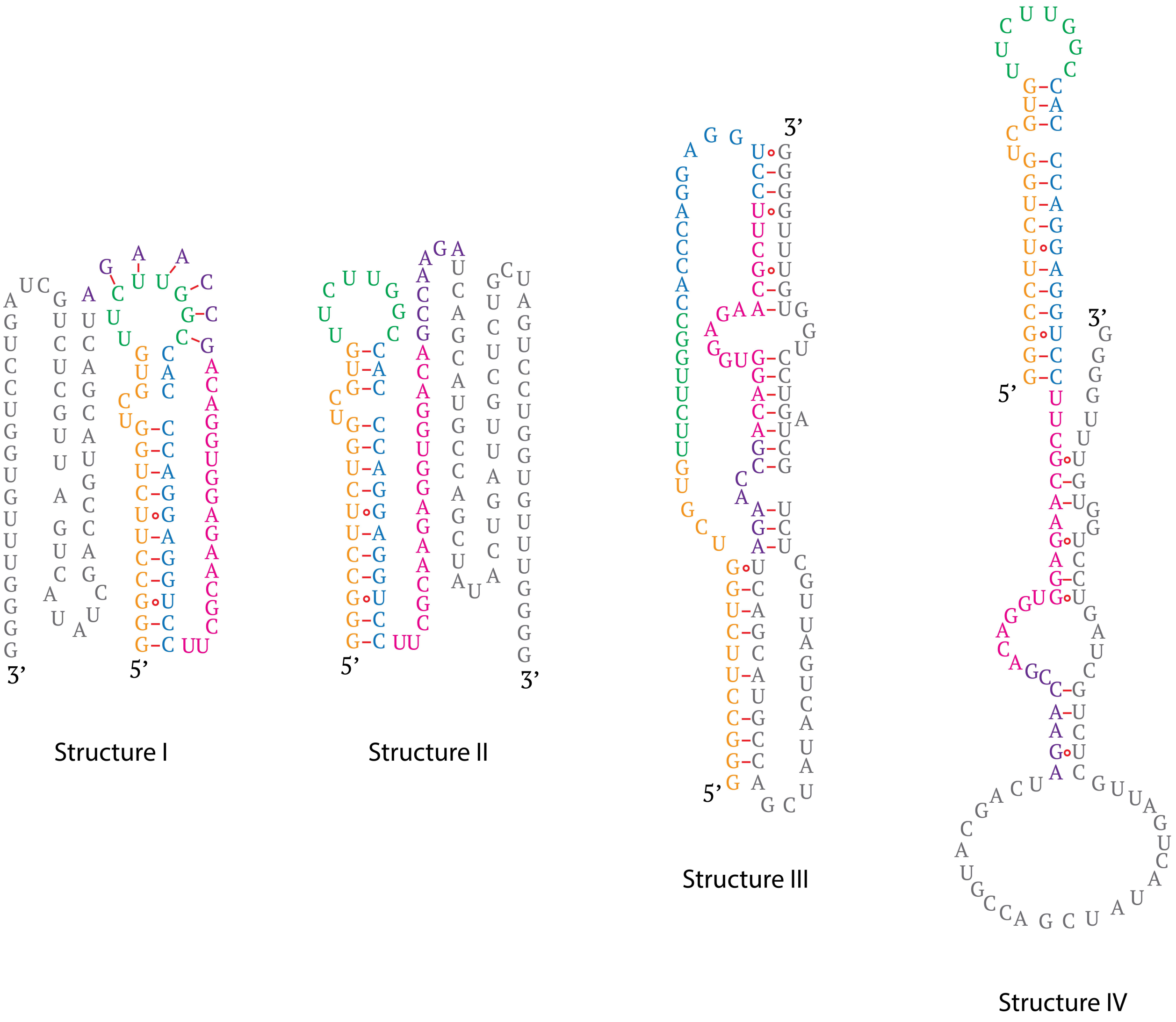


**Figure S4. Predicted secondary structures of the West Nile Virus (WNV) programmed -1 ribosomal frameshifting (PRF) site.** Color coding is used for the tracking of how different sequences are involved in the formation of the alternative secondary structures. Secondary structures were generated based on the data in (27).

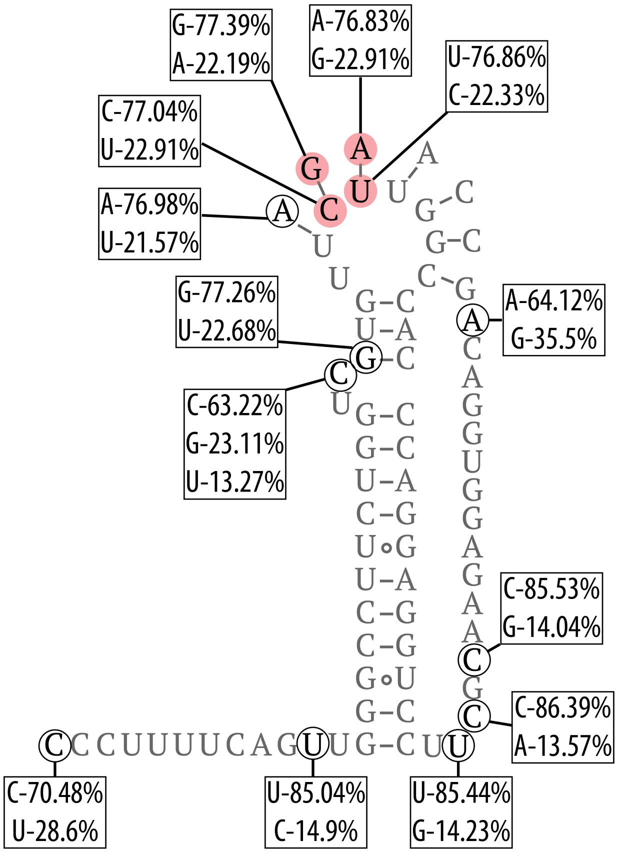


**Figure S5. Detailed analysis of nucleotide variation mapped onto the predicted secondary structure of the WNV pseudoknot.** The red-highlighted nucleotide pairs show concerted compensatory changes in the sequence consistent with the proposed secondary structure, though the statistical significance of these changes is low. These co-occurring changes support the WNV pseudoknot structure and are consistent with a conserved fold.

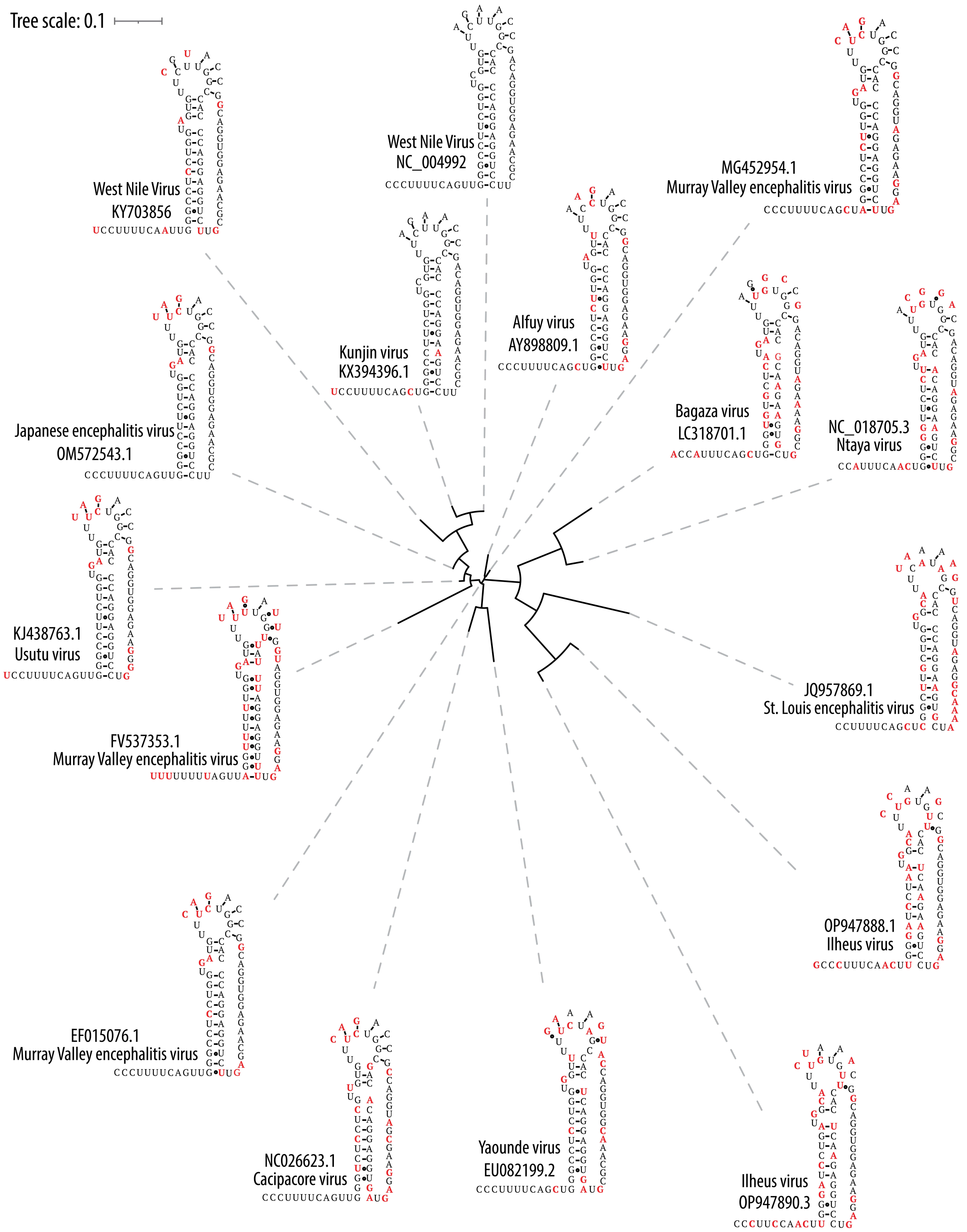


**Figure S6. Maximum-likelihood tree of *Flavivirus* frameshifting elements.** Maximum-likelihood tree of *Flavivirus* frameshifting elements with representative secondary structures. Sequence variations from the strain NY99 (RefSeq NC_009942) are colored in red.

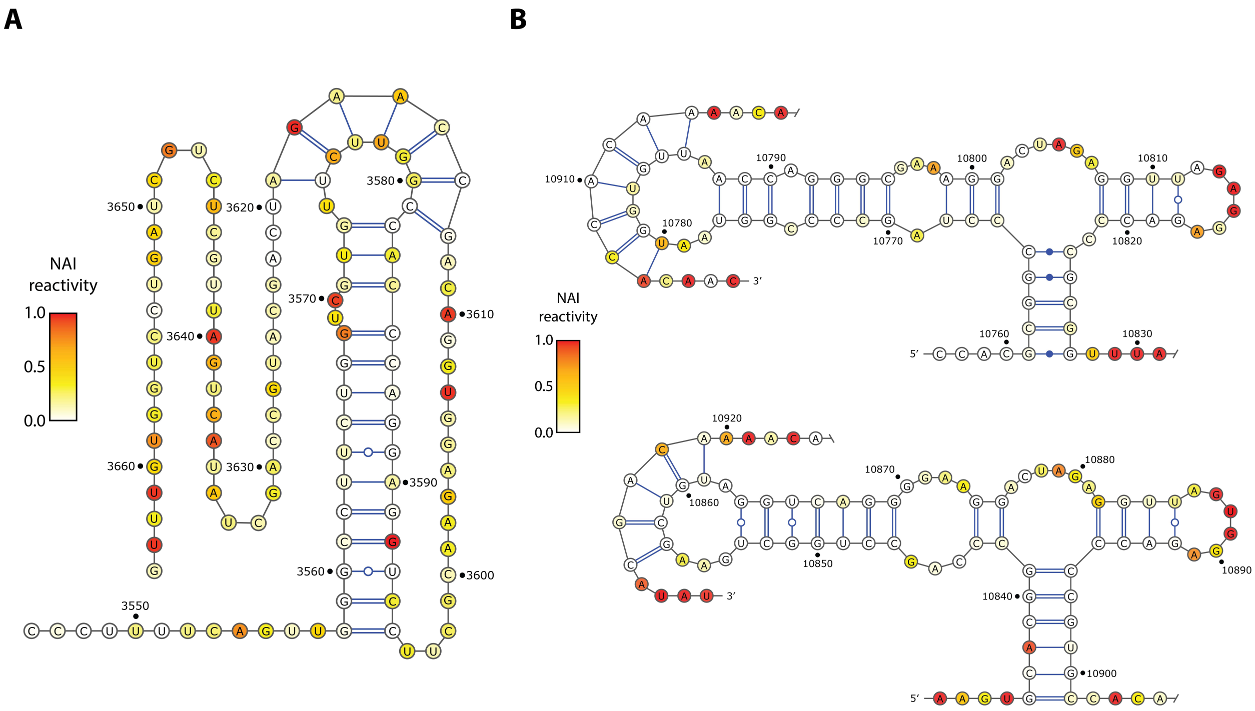


**Figure S7. Analysis of the predicted secondary structure and nucleotide reactivity of the WNV PRF element.**

(**A**) NAI reactivity profiling of the WNV PRF element from WNV-infected Vero cells mapped onto the proposed secondary structure. The downstream nucleotides exhibit elevated levels of NAI reactivity, indicating the absence of a stable secondary structure. The pseudoknot nucleotides show somewhat elevated reactivity levels, suggesting that the pseudoknot might possess conformational flexibility rather than being fully rigid.

(**B**) NAI reactivity profiling of two well-characterized dumbbell pseudoknot structures in the 3′ untranslated region (UTR) of WNV from WNV-infected Vero cells shown for comparison. These structures display much lower reactivity levels in known base-paired regions, reflecting the stability of their secondary structures in the same experiment. This comparison highlights how reactivity patterns likely correlate with structural stability in pseudoknots.

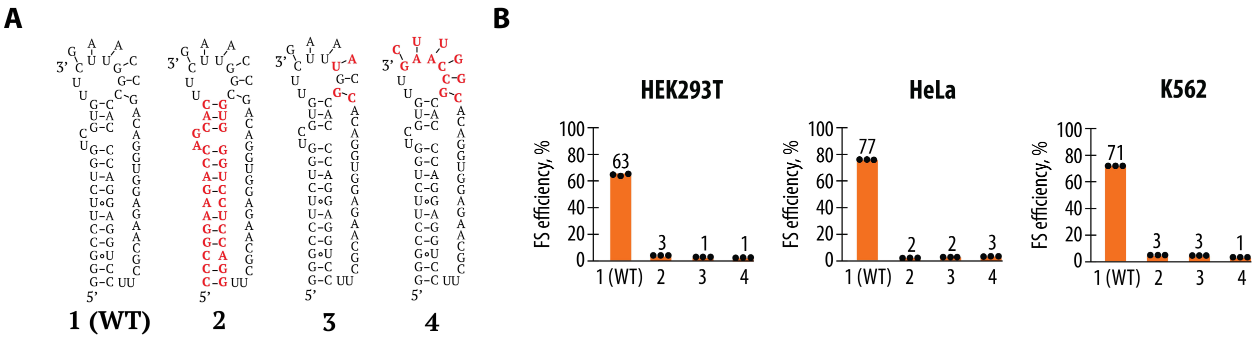


**Figure S8. Compensatory mutations aimed at restoring the predicted pseudoknot structure.** (**A**) Schematic representation of the compensatory mutations. (**B**) Effects of the compensatory mutations on frameshifting in HEK293T, HeLa, and K562 cells, compared to the wild-type (WT) construct. Bars represent the means of three independent experiments, with each dot corresponding to the individual experimental data points.

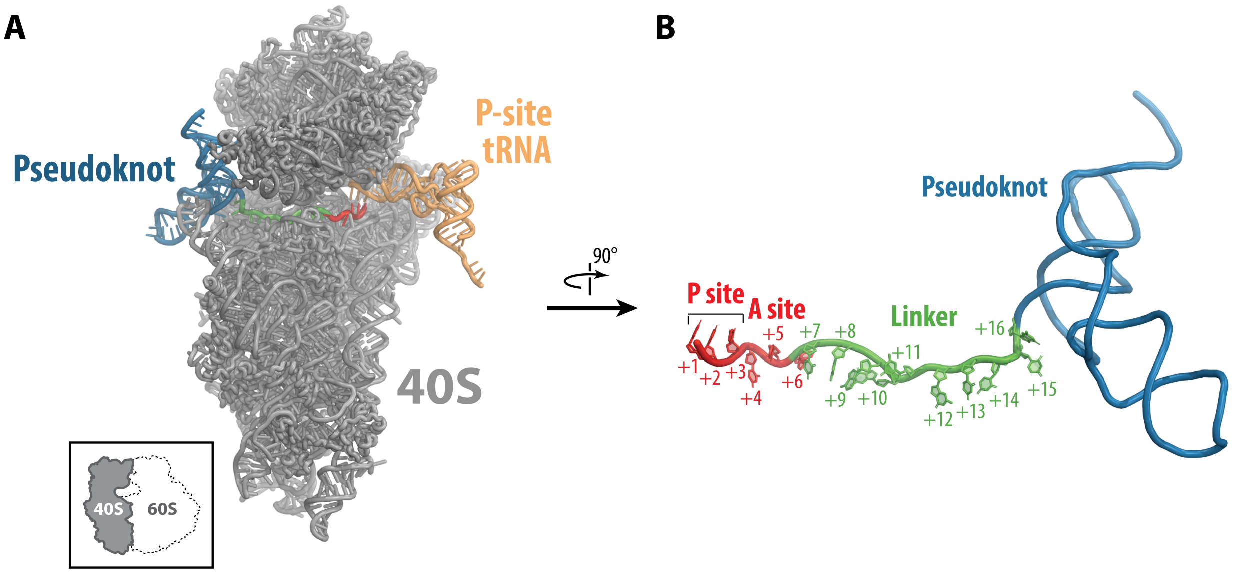


**Figure S9. Cryo-EM structure of the SARS-CoV-2 frameshifting site.** (**A**) Overview of the rabbit ribosome structure bound to mRNA containing the pseudoknot from SARS-CoV-2 (PDB ID: 7o7z). The 40S subunit is shown in gray, the P-site tRNA in orange, the pseudoknot in blue, the codons of the A- and P-sites in red, and the linker between the pseudoknot and the A- and P-site codons in green. The inset shows the 60S subunit removed for clarity, leaving only the 40S subunit shown in a side view. (**B**) Close-up view of the mRNA structure. The same color coding is used, with the 40S subunit not shown for clarity. The nucleotide numbering starts from the first nucleotide of the P-site codon (+1) and continues to the last unpaired nucleotide (+16). The view is from the solvent side of the 40S subunit in panel (**A**).

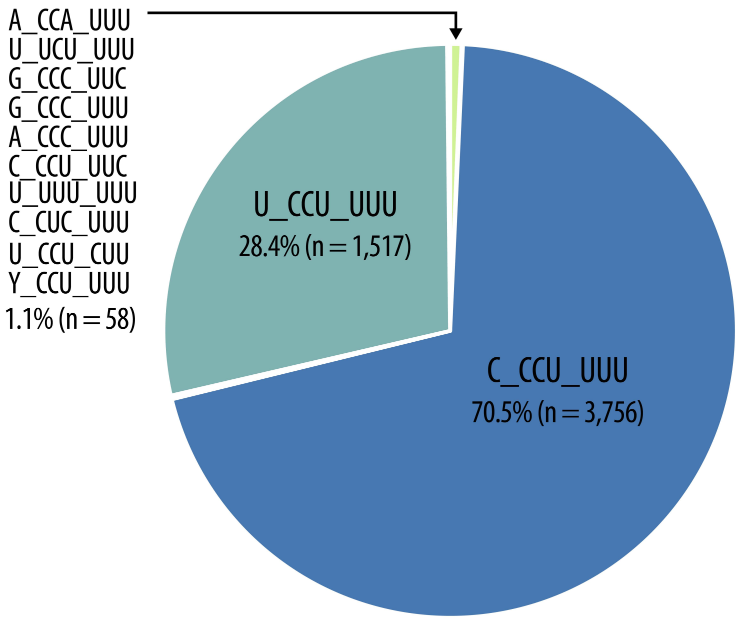


**Figure S10. Distribution of slippery sequences among flaviviruses.** The chart displays the proportion of different slippery sequences. The number of genomes and the corresponding percentages of the total are indicated for each sequence variant.

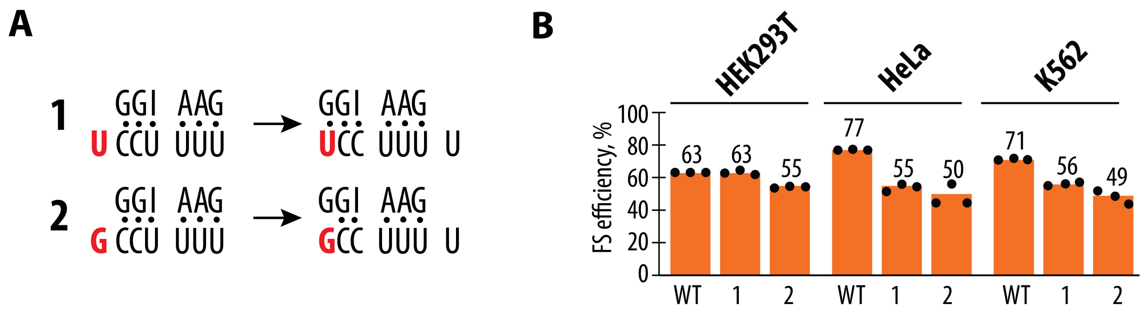


**Figure S11. Effect of the first nucleotide in the slippery site on frameshifting efficiency.** (**A**) Mutational analysis of the first nucleotide in the slippery site, with sequence register before (frame 0, left) and after (frame -1, right) frameshifting. (**B**) Frameshifting efficiency of constructs with mutations of the first nucleotide of slippery site, in HEK293T, HeLa, and K562 cells. Frameshifting efficiency is shown as a percentage of the ratio between normalized FLuc/RLuc luminescence in the -1 frame to the sum of normalized Fluc/Rluc luminescence in the -1 and 0 frames. Bars represent the means of three independent experiments, with each dot corresponding to the individual experimental data points.

**Supplementary Table S1**. Synthetic DNA blocks used in this study

| Figure | DNA name | Sequence |
|---|---|---|
| 1B | WT | TCGAGTCCAACCCCGGGCCCTACTCGAGCTTACCACCACTGCGCT<br>ACCAAACTGACAGCGGCTGTTGGTATGGTATGGAGATCAGACCAC<br>AGAGACATGATGAAAAGACCCTCGTGCAGTCACAAGTGAATGCT<br>TATAATGCTGATATGATTGACCCTTTTCAGTTGGGCCTTCTGGTCG<br>TGTTCTTGGCCACCCAGGAGGTCCTTCGCAAGAGGTGGACAGCCA<br>AGATCAGCATGCCAGCTATACTGATTGCTCTGCTAGTCCTGGTGTT<br>TGGGGGCATTACTTACGATCTGAGGCACGGCATAAGCAAAAGAT<br>CGTAG |
| | IFC | TCGAGTCCAACCCCGGGCCCTACTCGAGCTTACCACCACTGCGCT<br>ACCAAACTGACAGCGGCTGTTGGTATGGTATGGAGATCAGACCAC<br>AGAGACATGATGAAAAGACCCTCGTGCAGTCACAAGTGAATGCT<br>TATAATGCTGATATGATTGACCCTTTTCAGTTGGGCCTTCTGGTCG<br>TGTTCTTGGCCACCCAGGAGGTCCTTCGCAAGAGGTGGACAGCCA<br>AGATCAGCATGCCAGCTATACTGATTGCTCTGCTAGTCCTGGTGTT<br>TGGGGGCATTACTTACAGATCTGAGGCACGGCATAAGCAAAAGA<br>TCGTA |
| | SSM | TCGAGTCCAACCCCGGGCCCTACTCGAGCTTACCACCACTGCGCT<br>ACCAAACTGACAGCGGCTGTTGGTATGGTATGGAGATCAGACCAC<br>AGAGACATGATGAAAAGACCCTCGTGCAGTCACAAGTGAATGCT<br>TATAATGCTGATATGATTGATCGTACTCAGTTGGGCCTTCTGGTCG<br>TGTTCTTGGCCACCCAGGAGGTCCTTCGCAAGAGGTGGACAGCCA<br>AGATCAGCATGCCAGCTATACTGATTGCTCTGCTAGTCCTGGTGTT<br>TGGGGGCATTACTTACGATCTGAGGCACGGCATAAGCAAAAGAT<br>CGTAG |
| | 5’ STOP | TCGAGTCCAACCCCGGGCCCTACTCGAGCTAATTACCACCACTGC<br>GCTACCAAACTGACAGCGGCTGTTGGTATGGTATGGAGATCAGAC<br>CACAGAGACATGATGAAAAGACCCTCGTGCAGTCACAAGTGAAT<br>GCTTATAATGCTGATATGATTGACCCTTTTCAGTTGGGCCTTCTGG<br>TCGTGTTCTTGGCCACCCAGGAGGTCCTTCGCAAGAGGTGGACAG<br>CCAAGATCAGCATGCCAGCTATACTGATTGCTCTGCTAGTCCTGG<br>TGTTTGGGGGCATTACTTACGATCTGAGGCACGGCATAAGCAAAA<br>GATCG |
| 2A | Mutant NC WT | GGCGGGAGACGTCGAGTCCAACCCCGGGCCCTACTCGAGCACCA<br>CCACTGCGCTACCAAACCGACAGCGGCTGTTGGTATGGTATGGAG<br>ATCAGACCACAGAGACACGACGAAAAGACCCTCGTGCAGTCACA<br>AGTGAATGCTTACAATGCCGATATGATTTTGACCCTTTTCAGTTGG<br>GCCTTCTGGTCGTGTTCTTGGCCACCCAGGAGGTCCTTCGCAAGA<br>GGTGGACAGCCAAGATCAGCATGCCAGCTATACTGATTGCTCTGC |

| | | |
|---|---|---|
| | | TAGTCCTGGTGTTTGGGGGCATTACTTACGATCTGAGGCACGGCATAAGCAAAAGATCGTAGCCCCAGTAAAGCAAACACTCAACTTCGATCTACTCAAACTCGCAGGTGATGTGGAATCTAATCCAGGACCTTTCGGATCTGCCACCATTGAAGATGCCAAAAACATTAAGAAGGGCCCAGCGCCATTCTACCCACTCGAAGACGG |
| | Mutant NC IFC | GGCGGGAGACGTCGAGTCCAACCCCGGGCCCTACTCGAGCACCACCACTGCGCTACCAAACCGACAGCGGCTGTTGGTATGGTATGGAGATCAGACCACAGAGACACGACGAAAAGACCCTCGTGCAGTCACAAGTGAATGCTTACAATGCCGATATGATTTTGACCCTTTTCAGTTGGGCCTTCTGGTCGTGTTCTTGGCCACCCAGGAGGTCCTTCGCAAGAGGTGGACAGCCAAGATCAGCATGCCAGCTATACTGATTGCTCTGCTAGTCCTGGTGTTTGGGGGCATTACTTACAGATCTGAGGCACGGCATAAGCAAAAGATCGTAGCCCCAGTAAAGCAAACACTCAACTTCGATCTACTCAAACTCGCAGGTGATGTGGAATCTAATCCAGGACCTTTCGGATCTGCCACCATTGAAGATGCCAAAAACATTAAGAAGGGCCCAGCGCCATTCTACCCACTCGAAGACGG |
| 5B | -2 reporter WT | GGCGGGAGACGTCGAGTCCAACCCCGGGCCCTACTCGAGCTTACCACCACTGCGCTACCAAACTGACAGCGGCTGTTGGTATGGTATGGAGATCAGACCACAGAGACATGATGAAAAGACCCTCGTGCAGTCACAAGTGAATGCTTATAATGCTGATATGATTGACCCTTTTCAGTTGGGCCTTCTGGTCGTGTTCTTGGCCACCCAGGAGGTCCTTCGCAAGAGGTGGACAGCCAAGATCAGCATGCCAGCTATACGCACGGCATAAGGTAAAGCAAACACTCAACTTCGATCTACTCAAACTCGCAGGTGATGTGGAATCTAATCCAGGACCTTTCGGATCTGCCACCATTGAAGATGCCAAAAACATTAAGAAGGGCCCAGCGCCATTCTACCCACTCGAAGACGG |
| | -2 reporter IFC | GGCGGGAGACGTCGAGTCCAACCCCGGGCCCTACTCGAGCTTACCACCACTGCGCTACCAAACTGACAGCGGCTGTTGGTATGGTATGGAGATCAGACCACAGAGACATGATGAAAAGACCCTCGTGCAGTCACAAGTGAATGCTTATAATGCTGATATGATTGACCCTTTTCAGTTGGGCCTTCTGGTCGTGTTCTTGGCCACCCAGGAGGTCCTTCGCAAGAGGTGGACAGCCAAGATCAGCATGCCAGCTATACGCACGGCAGTAAAGCAAACACTCAACTTCGATCTACTCAAACTCGCAGGTGATGTGGAATCTAATCCAGGACCTTTCGGATCTGCCACCATTGAAGATGCCAAAAACATTAAGAAGGGCCCAGCGCCATTCTACCCACTCGAAGACGG |

**Supplementary Table S2.** Primers used in this study

| Figure | Primer name | Sequence |
|---|---|---|
| 1B | 3' control PCR Fwd | TACGATCTTTTGCTTATGCCGTGCC |
| | 3' control PCR Rev | AAGCAAACACTCAACTTCGATCTAC |
| | 3' control Gibson insert | GAGGCACGGCATAAGCAAAAGATCGTAGCC TAAGTAAAGCAAACACTCAACTTCGATCTA |
| 1D | Vector PCR Fwd | ACCATACCCATACGACGTACCAGATTACGCTTAGCGT CTTCACACTCGAAGATTTCGTTG |
| | Vector PCR Rev | CAGTTTGGTAGCGCAGCAGTGGTAAGCCACCAGAGCCC ATATGCCCTCCA |
| | Insert PCR Fwd | TGGAGGGCATATGGGCTCTGGTGGCTTACCACCACTGCGC ATCCAACTGCGCTACCAAACTG |
| | Insert PCR Rev | GCTAAGCGTAATCTGGTACGTCGTATGGGTATGGTAAGTAA TGCCCCCAAACACCAGG |
| 2D | 5' truncation PCR FWD | GCTCGAGTAGGGCCC |
| | 5' truncation PCR Rev | GACCCTTTTCAGTTGGGC |
| | 3' truncation WT PCR FWD | GTATAGCTGGCATGCTG |
| | 3' truncation WT PCR Rev | GATCTGAGGCACGGC |
| | 3' truncation IFC PCR FWD | GTATAGCTGGCATGCTG |
| | 3' truncation IFC PCR Rev | AGATCTGAGGCACGGC |

| | | |
|---|---|---|
| | Short insert PCR WT1 | GAGTCCAACCCCGGGCCCTACTCGAGCGACCCTTTTCAGTTGGGC CTTCTGGTCGTGTTC |
| | Short insert PCR WT2 | CAGTTGGGCCTTCTGGTCGTGTTCTTGGCCACCCAGGAGGTC CTTCGCAAGAGGTGGAC |
| | Short insert PCR WT3 | TGCTTATGCCGTGCCTCAGATCGCTGATCTTGGCTGTCCACC TCTTGCGAAGGACCTCCT |
| | Short insert PCR IFC1 | GAGTCCAACCCCGGGCCCTACTCGAGCGACCCTTTTCAGT TGGGCCTTCTGGTCGTGTTC |
| | Short insert PCR IFC2 | CAGTTGGGCCTTCTGGTCGTGTTCTTGGCCACCCAGGAGGTC CTTCGCAAGAGGTGGAC |
| | Short insert PCR IFC3 | TGCTTATGCCGTGCCTCAGATCTGCTGATCTTGGCTGTCCACC TCTTGCGAAGGACCTCC |
| 3A | Construct 2 Fwd | ACCAGCACTTCTTGGCCACCCAG |
| | Construct 2 Rev | CTTCCGGGAACTGAAAAGGGTCAATCATATC |
| | Construct 3 Fwd | CTCCAGGTTCGCAAGAGGTGGAC |
| | Construct 3 Rev | GACCCACGCCAAGAACACGACC |
| | Construct 4 Fwd | TCCACCTGTGCCAAGATCAGCATGC |
| | Construct 4 Rev | GAACGCTTGGACCTCCTGGGTGGG |
| | Construct 5 Fwd | GTGTTCTTTGGCACCCAGGAGGTCCTTC |
| | Construct 5 Rev | GACCAGAAGGCCCAACTGAAAAGGGTC |
| | Construct 6 Fwd | TGGTCGTGTTGAACCHCACCCAGGAG |

| | | |
|---|---|---|
| | Construct 6 Rev | GAAGGCCCAACTGAAAAG |
| 4A | 2 nt linker PCR Fwd | GGGCCTTCTGGTCGTG |
| | 2 nt linker PCR Rev | TGAAAAGGGTCAATCATATCAGC |
| | 8 nt linker PCR Fwd | TTTGGGCCTTCTGGTCGTG |
| | 8 nt linker PCR Rev | AACTGAAAAGGGTCAATCATATCAG |
| | 11 nt linker PCR Fwd | TTTAAAGGGCCTTCTGGTCGTG |
| | 11 nt linker PCR Rev | AACTGAAAAGGGTCAATCATATCAG |
| 4C | Construct 2 PCR Fwd | GAAGGCGGAACTGAAAAGGGTCAATCATATC |
| | Construct 2 PCR Rev | TGGTCGTGTTCTTGGCCACCCAG |
| | Construct 3 PCR Fwd | GGTCGTGTTCTTGGCCACCCAGGAG |
| | Construct 3 PCR Rev | AGAAGGGGGAACYGAAAAGGGTCAATCATATC |
| | Construct 4 PCR Fwd | GGTCGTGTTCTTGGCCACCCAGGAG |
| | Construct 4 PCR Rev | AGAAGCGGGAACTGAAAAGGGTCAATCATATC |
| | Construct 5 PCR Fwd | TGGTCGTGTTCTTGGCCACCCAG |
| | Construct 5 PCR Rev | GAACCGGGAACTGAAAAGGGTCAATCATATC |
| | Construct 6 PCR Fwd | TGGTCGTGTTCTTGGCCACCCAG |

| | | |
|---|---|---|
| | Construct 6 PCR Rev | CTTCCGGGAACTGAAAAGGGTCAATCATATC |
| 5B | U_UUU_UUU slippery site PCR Fwd | TTTTCAGTTGGGCCTTCTG |
| | U_UUU_UUU slippery site PCR Rev | AAATCAATCATATCAGCATTATAAGC |
| 6B | P1 Fwd | TTTTCAGTTGGGCCTTCTG |
| | P1 Rev | GCGTCAATCATATCAGCATTATAAGC |
| | P2 Fwd | TTTTCAGTTGGGCCTTCTG |
| | P2 Rev | CGGTCAATCATATCAGCATTATAAGC |
| | P3 Fwd | GGGTCAATCATATCAGCATTATAAGC |
| | P3 Rev | GTTTCAGTTGGGCCTTCTG |
| | P4 Fwd | ATTTCAGTTGGGCCTTCTG |
| | P4 Rev | GGGTCAATCATATCAGCATTATAAGC |
| | P5 Fwd | CTTTCAGTTGGGCCTTCTG |
| | P5 Rev | GGGTCAATCATATCAGCATTATTAGC |
| 6D | A1 Fwd | AAAACAGTTGGGCCTTCTG |
| | A1 Rev | GGGTCAATCATATCAGCATTATAAGC |
| | A2 Fwd | TTTCCAGTTGGGCCTTCTG |
| | A2 Rev | GGGTCAATCATATCAGCATTATAAGC |
| | A3 Fwd | TGTTCAGTTGGGCCTTCTG |
| | A3 Rev | GGGTCAATCATATCAGCATTATAAGC |
| | A4 Fwd | TTGTCAGTTGGGCCTTCTG |
| | A4 Rev | GGGTCAATCATATCAGCATTATAAGC |
| | A5 Fwd | TTTGCAGTTGGGCCTTCTG |
| | A5 Rev | GGGTCAATCATATCAGCATTATAAGC |

| | | |
|---|---|---|
| S8 | Construct 2 PCR FWD | TTGGCGTGGGTCCTCCAGGTTCGCAAGAGGTGGAC |
| | Construct 2 PCR Rev | GAAGTGCTGGTCTTCCGGGAACTGAAAAGGGTCAATCATATC |
| | Construct 3 PCR FWD | GTGTTCTTTGGCACCCAGGAGGTCCTTC |
| | Construct 3 PCR Rev | GACCAGAAGGCCCAACTGAAAAGGGTC |
| | Construct 4 PCR FWD | CGGTTCATCAGCATGCCAGCTATAC |
| | Construct 4 PCR Rev | TGTCCACCTCTTGCGAAGGAC |
| S10 | Construct 1 PCR FWD | TTTTCAGTTGGGCCTTCTG |
| | Construct 1 PCR Rev | GGATCAATCATATCAGCATTATAAGC |
| | Construct 2 PCR FWD | TTTTCAGTTGGGCCTTCTG |
| | Construct 2 PCR Rev | GGCTCAATCATATCAGCATTATAAGC |
